# Differential control of growth and identity by HNF4α isoforms in pancreatic ductal adenocarcinoma

**DOI:** 10.1101/2025.03.31.646428

**Authors:** Pengshu Fang, Emily Wilson, Chris Stubben, Acramul Kabir, Kajsa Affolter, Xiaoyang Zhang, Eric L. Snyder

## Abstract

**Background:** Although transcriptomic studies have stratified pancreatic ductal adenocarcinoma (PDAC) into clinically relevant subtypes, classical or basal-like, further research is needed to identify the transcriptional regulators of each subtype. Previous studies identified HNF4α as a key regulator of the classical subtype, but the distinct contributions of its isoforms (P1 and P2), which display dichotomous functions in normal development and gastrointestinal malignancies, remain unexplored.

**Objective:** The objective of this study is to investigate the role of HNF4α P1 and P2 isoforms in regulating growth and differentiation.

**Design:** We performed functional, transcriptomic, and epigenetic analysis after exogenous expression in HNF4α-negative models or CRISPRi-mediated knockdown of endogenous isoforms.

**Results:** We characterized the variable expression of P1 isoforms in HNF4α-positive tumors. We demonstrate that P1 isoforms are less compatible with growth than P2 isoforms. Despite sharing a common DNA binding domain, we show that P1 isoforms are stronger transcriptional regulators.

**Conclusions:** Our study characterizes the functional roles of HNF4α P1 and P2 isoforms in PDAC and highlights the necessity of considering different isoforms when studying molecular regulators.

## INTRODUCTION

Despite its low incidence rate, pancreatic cancer is the 4^th^ leading cause of cancer death in the United States each year^1^. The high mortality rate of pancreatic cancer, of which pancreatic ductal adenocarcinoma (PDAC) composes >90% of cases, is because it is often diagnosed at an advanced stage and is refractory to standard of care treatments^2^. While most tumors harbor *KRAS* (>90%) and *TP53* (>70%) mutations, transcriptomic studies within the last decade have identified clinically relevant molecular subtypes of PDAC^3–5^. PDAC can be classified into a classical, pancreatic progenitor subtype or a basal-like, squamous subtype^6^. The classical subtype is associated with better survival and response to chemotherapy^4,7,8^, lower grade tumors^9^, and higher expression of endodermal lineage specifiers (e.g., *HNF4A, GATA6*)^5,7^ than the basal-like subtype, which has a greater sensitivity to targeted therapy including KRAS inhibition^8,10^. Despite this binary classification’s robustness and clinical relevance, there is evidence for heterogeneity within these broader subtypes that is not yet fully understood. A recent study has further stratified the classical subtype into classical-A (gastric) and classical-B (intestinal)^11^. Thus, more research is needed to identify the precise molecular regulators of this intra-subtype heterogeneity.

We and others have shown that HNF4α is a critical regulator of molecular subtype, growth, and metabolism in PDAC^12–14^. HNF4α is a nuclear hormone receptor that is essential for the development and maintenance of the gastrointestinal (GI) tract^15–17^ and cellular and xenobiotic metabolism^18,19^. We demonstrated that loss of HNF4α at tumor initiation in a KRAS-driven, loss-of-p53, genetically engineered mouse model (GEMM) led to an increase in tumor burden and a decrease in overall survival^12^. Integrated RNAseq/ChIP-seq analysis revealed that HNF4α binds and activates genes associated with the classical subtype. HNF4α also represses a subset of genes enriched in basal-like PDAC (e.g., *SIX4*), but loss of HNF4α was insufficient for complete conversion to the basal-like subtype^12^. Consistent with this, another study demonstrated that concomitant loss of GATA6, HNF4α, and HNF1α is required to activate the basal-like program^14^. Lastly, HNF4α regulates metabolism within the classical subtype, as HNF4α knockdown resulted in a switch from dependence on oxidative phosphorylation to glycolysis^13^.

There are 12 isoforms of HNF4α driven by expression from two promoters, P1 and P2^20–23^. In the GEMM, HNF4α-positive PDAC only express P2 isoforms^12^. In contrast, we observed that HNF4α-positive patient-derived xenografts (PDXs) were uniformly P2-positive but expressed variable levels of P1 isoforms, indicating GEMMs do not fully recapitulate the heterogeneity found in human PDAC^12^. P1 and P2 isoforms contain unique first exons but are then spliced into shared DNA and ligand binding domains. Exon 1 of P1 isoforms contains an AF-1 transactivation domain, whereas P2 isoforms do not, allowing P1 isoforms to interact with additional cofactors^21^. HNF4α isoforms display tissue-specific expression patterns in the GI tract; the stomach and pancreas predominantly express P2, the adult liver and kidney express P1, and the intestines express both isoforms^24^. In general, proliferative cells (e.g., fetal liver and colonic crypts) express P2 isoforms, whereas P1 isoforms are expressed in differentiated cells (e.g., adult liver and colonocytes)^25–27^. Although the role of each isoform in PDAC is unknown, P1 isoforms restrain tumor growth, while P2 isoforms are permissive for tumor growth in other GI malignancies (colorectal and hepatocellular carcinoma^26–28^). These observations suggest that distinct isoforms of HNF4α may play unique roles in PDAC and regulate heterogeneity within classical tumors.

Previously, we demonstrated that HNF4α is a key regulator of growth and differentiation in PDAC and observed heterogeneous expression of P1 and P2 isoforms in human PDXs^12^. Based on the expression patterns and function of HNF4α isoforms in normal GI tissues and other GI malignancies, we investigated the roles of HNF4α isoforms in PDAC. In both our gain-of-function and knockdown studies, we found that P1 isoforms were less compatible with growth and more potent transcriptional regulators than P2 isoforms. While P1 and P2 isoforms did not promote unique GI lineages, P1 isoforms pushed cells to a further differentiated state along similar lineage trajectories.

## RESULTS

### Exogenous P1 isoforms restrain growth in vitro to a greater degree than P2 isoforms

We first used isoform-specific antibodies to evaluate the expression of HNF4α-P1 and P2 on a panel of primary human PDAC. In tumors with diffuse positivity for total HNF4α (n=9), we found that P2 was expressed in >90% of carcinoma cells. In contrast, the percentage of P1-positive cells was highly variable (range: 12-63%) and lower than P2 in any tumor (**Figure 1A-B, Supplemental Table S1**). These data are consistent with our prior analysis of human PDXs^12^ and demonstrate that there is significant intra– and inter-tumoral heterogeneity of P1 expression in human PDAC. Of note we did not observe a correlation between P1 positivity and any specific morphologic features.

**Figure 1:**
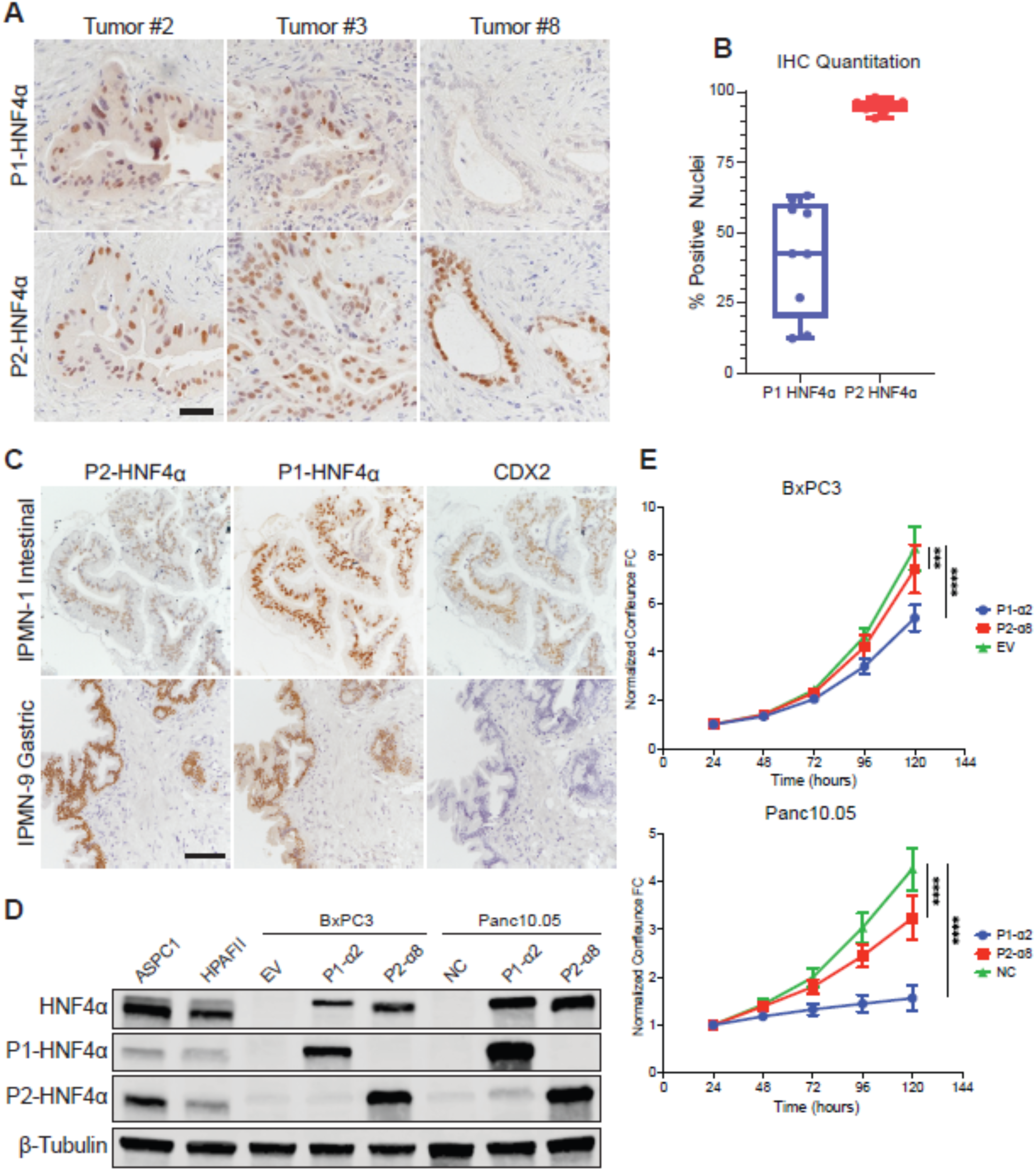
Exogenous expression of P1-HNF4α restrains tumor growth in vitro. **A.** Representative IHC for HNF4α P1 and P2 isoforms in human PDAC tumors (scale=100 microns). **B.** Percentage of PDAC nuclei positive for HNF4α P1 or P2 isoforms (n=9), only neoplastic cells were included in the quantitation while stromal cells were omitted. **C.** Representative IHC for HNF4α P1 and P2 isoforms and CDX2 in human IPMN neoplasia (scale=250 microns). **D.** Representative immunoblot of total HNF4α, P1-HNF4α, P2-HNF4α, or β-Tubulin in BxPC3 and Panc10.05 cells treated with dox to induce expression of P1-α2 or P2α8. ASPC1 and HPAFII express endogenous HNF4α and are positive controls. Empty vector (BxPC3) or GFP (Panc10.05) are negative controls (NC). **E.** Incucyte growth assay in BxPC3 and Panc10.05 after dox induction of negative control, P1-α2, or P1-α8 for 5 days. Confluence was normalized to the initial seeding density at 24 hours, 8 replicates, 3 independent experiments per cell line. Mean +/-SEM shown. Significance was calculated by Log_2_ transformation of the data and testing the differences in the slope via an ANCOVA test (***=p<.001, ****=p<.0001, **Supplemental** Figure 1F).

Intraductal papillary mucinous neoplasms (IPMNs) are macroscopic PDAC precursor lesions that can be subclassified by their resemblance to normal GI tissues (e.g., foveolar/gastric, intestinal, or pancreatobiliary). A recent study demonstrated that P1 isoforms exhibit a distinct spatial distribution in high-grade IPMN that mimics the normal intestine^29^. However, this study did not directly assess whether isoform expression in gastric vs. intestinal IPMNs mimics expression in normal tissues. We therefore performed IHC for P1, P2, and intestinal marker CDX2 on a panel of 8 IPMN cases (3 intestinal and 5 gastric). P2 was uniformly expressed in both subtypes, consistent with Wong et al. and our observations in PDAC. As expected, all three intestinal-type IPMNs expressed P1 and CDX2 in a subset of neoplastic cells, whereas CDX2 was negative in gastric-type IPMNs. Surprisingly, all five gastric-type IPMNs expressed P1 in a subset of neoplastic cells, in contrast with the P2-only phenotype of normal stomach (**Figure 1C, Supplemental Table S1**). Taken together, these data show that P1 expression can become uncoupled from normal GI differentiation programs early in the progression of PDAC neoplasia.

To investigate the relative ability of HNF4α isoforms to regulate PDAC growth in vitro, we expressed exogenous, doxycycline-inducible (dox), HNF4α2 and HNF4α8 as representative P1 and P2 isoforms, respectively (**Supplemental Figure S1A**) in the HNF4α-negative cell lines BxPC3 (squamous-like, ΔNP63-positive) and Panc10.05 (poorly differentiated). HNF4α2 and HNF4α8 are two of the primary isoforms expressed from the P1 and P2 promoter^26^, respectively, and both have been previously used as representative HNF4α isoforms^28,30^. Dox treatment induced near equivalent levels of P1 and P2 isoforms within each HNF4α-negative cell line that were within the range of total endogenous HNF4α expression in two PDAC cell lines (ASPC1 and HPAFII, **Figure 1D**). Overall levels of P1-α2 and P2-α8 in Panc10.05 were higher than in BxPC3 (**Figure 1D**) despite using 20-fold less dox in Panc10.05 (25ng/mL vs 500ng/mL). Using Incucyte assays, we found that exogenous P1-α2 restrained growth to a greater extent than P2-α8 in both cell lines (**Figure 1E and Supplemental Figure S1B**).

### Exogenous P1 isoforms are stronger transcriptional regulators than P2 isoforms

Due to the differential effects of P1-α2 and P2-α8 in regulating growth, we sought to investigate their transcriptional activity. We performed RNAseq on BxPC3 (**Figure 2**) and Panc10.05 (**Supplemental Figure S2**) after exogenous expression of isoforms. Subsequently, we performed DESeq2 on P1-α2 vs. control, P2-α8 vs. control, and P1-α2 vs. P2-α8 to determine the number of differentially expressed genes (DEGs). In BxPC3, expression of P1-α2 led to a greater number of significant DEGs (542 upregulated, 198 downregulated) than P2-a8 (228 upregulated, 54 downregulated) vs. control (Padj<.05, |log2FC|>1, **Supplemental Table S2**). We performed Gene Set Enrichment Analysis (GSEA) (**Figure 2A**) and observed that both P1 and P2 isoforms were enriched for intestinal and hepatic signatures, classical PDAC subtypes, and metabolism signatures. However, when contrasting P1 vs. P2 to isolate genes more strongly regulated by one isoform, all of the above signatures were enriched in P1. Additionally, we observed that P1-α2 inhibited signatures of both normal squamous cell types and the basal-like PDAC subtypes to a greater degree than P2-α8. Lastly, we observed a negative enrichment for E2F and G2M signatures, specifically in P1-α2, in concordance with our in vitro growth assay (**Figure 1E**). Based on the spatial expression of HNF4α isoforms in the colon, we asked whether P1 and P2 isoforms were differentially enriched for intestinal cell types. We performed GSEA analysis of HNF4α-dependent intestinal signatures^31^ and observed that while both P1-α2 and P2-α8 had similar enrichment for differentiated cell and enterocyte signatures, P1-α2 were more negatively enriched for a proliferative cell signature (**Figure 2B**, –2.6 NES vs –1.5 NES). This negative enrichment may explain the greater inhibitory effect of P1-α2 in our incucyte assay (**Figure 1E**).

**Figure 2:**
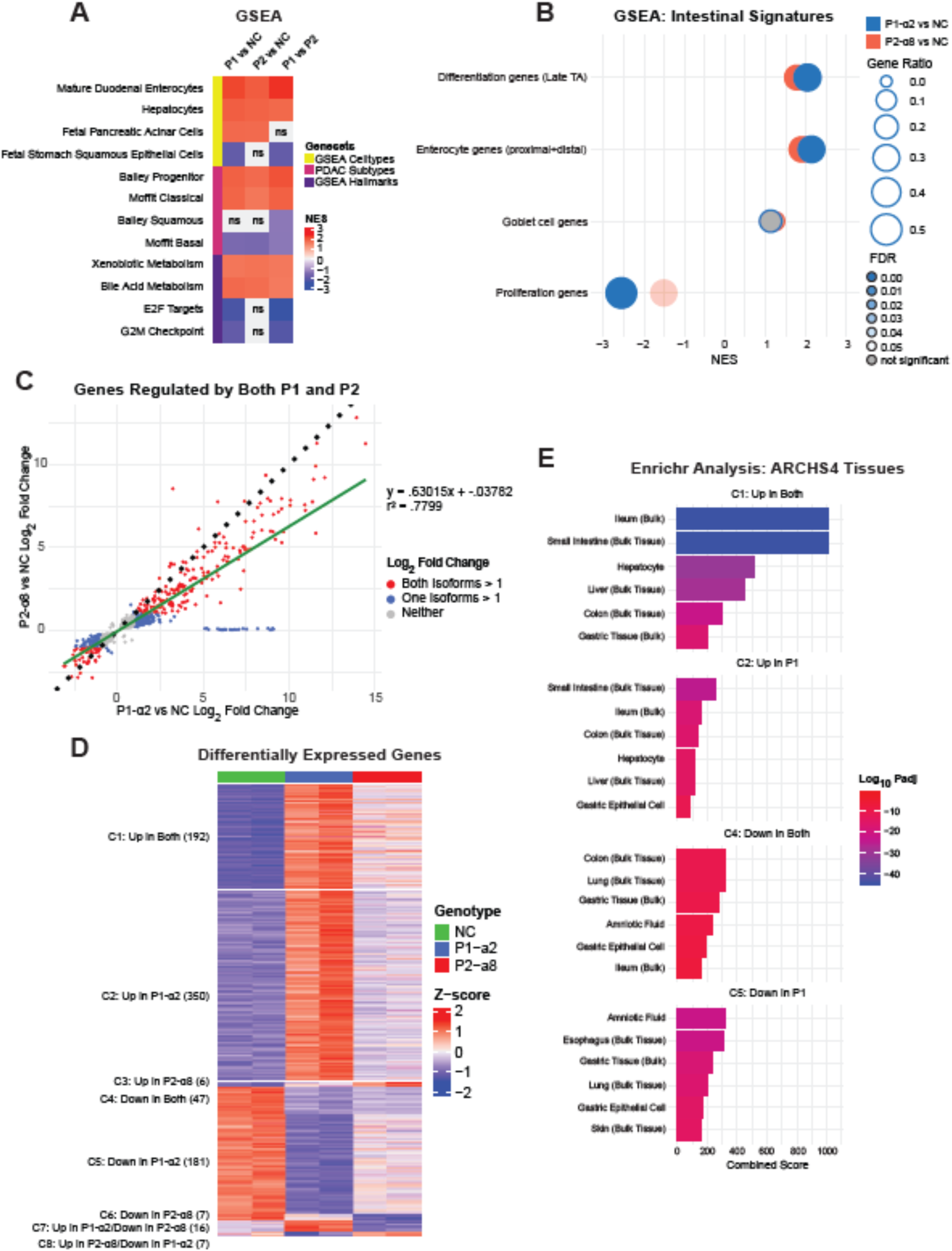
Exogenous P1-HNF4α is a stronger transcriptional regulator than P2-HNF4α in BxPC3. RNAseq analysis following expression of P1-HNF4α2 or P2-HNF4α8 in BxPC3. **A**. Heatmap of GSEA results for the hallmark, cell type, and PDAC subtype gene sets in the following contrasts: P1-α2 vs. NC, P2-α8 vs. NC, and P1-α2 vs. P2-α8. In the P1-α2 vs. P2-α8 contrast, a positive NES score indicates enrichment in P1, whereas a negative NES score indicates enrichment in P2. GSEA results were significant in at least one contrast (FDR<.05), ns=not significant. **B**. Dotplot of GSEA results for intestinal signatures (converted from mouse to human genes) used in Chen et al. 2019. **C**. Scatterplot of the log_2_ fold change of DEGs significantly regulated by both isoforms (Padj<.05). Green-line represents a linear regression model of the data (y=.63015x+-.03782, r^2^=.7799), dotted-line represents a slope of 1. **D**. Heatmap of manually clustered DEGs significantly regulated in at least one contrast (|log2FC|>1, Padj<.05). **E**. Enrichr analysis results for the indicated clusters in the ARCHS4 Tissues gene set.

We then plotted the log2FC of genes significantly regulated by both isoforms (Padj<.05) to determine which isoform was the stronger transcriptional regulator of these shared target genes (n=952, **Figure 2C**). We fit a linear model of the data, which yielded a slope of .63, demonstrating that P1 isoforms are stronger transcriptional regulators of most shared target genes. Of note, the magnitude of gene activation was greater than the magnitude of repression, suggesting HNF4α primarily acts as a transcriptional activator. Lastly, we clustered and plotted a heatmap of all significant DEGs from each contrast based on the directionality of gene expression (Padj<.05, |log2FC|>1, **Figure 2D, Supplemental Table S3**), resulting in 8 clusters. In clusters C1 (induced by both isoforms) and C4 (inhibited by both isoforms), P1-α2 expression led to a higher and lower z-score than P2-α8, respectively, further demonstrating that P1-α2 is a stronger transcriptional regulator of genes regulated by both isoforms. When examining clusters of genes uniquely regulated by one isoform (C2 (up in P1), C3 (up in P2), C6 (down in P1), and C6 (down in P2)), we found that P1 isoforms also regulated a greater absolute number of DEGs than P2 isoforms (350 vs 6 upregulated and 181 vs 7 downregulated, respectively).

We performed Enrichr analysis for the ARCHS4 Tissues gene sets for clusters 1, 2, 4, and 5 to identify cell type signatures enriched in each cluster (**Figure 2E, Supplemental Table S3**). We found that C1 genes exhibited the greatest tissue specificity (based on score and P value) and were highly enriched for intestinal/liver signatures, while C2 was also enriched for these signatures, albeit with a lower score. A signature of normal squamous epithelium (esophagus) was the most enriched in C5 (repressed by P1), consistent with the ability of P1 to inhibit squamous/basal-like PDAC signatures to a greater extent than P2 (**Figure 2A**). Analysis of C4 genes (repressed by both isoforms) did not yield a coherent pattern, potentially due to the small number of genes. Although both clusters of repressed genes (C4 and C5) have some enrichment for gastric signatures, further examination revealed that the predominant genes driving this enrichment are broadly expressed in multiple tissue types, including gastric. Taken together, these data show that both isoforms activate canonical intestinal/hepatic identity programs, but P1 is a stronger inducer of these identities due to stronger activation of common targets, upregulation of unique genes found in fully differentiated cells, and stronger inhibition of non-GI signatures.

We performed HNF4α ChIP-seq to investigate whether the change in DEGs after dox-induced expression of P1 and P2 isoforms resulted from direct DNA binding using a pan-HNF4α antibody (**Figure 3**). Although expression levels of P1 and P2 isoforms were similar (**Figure 1C**), P1-α2 bound a greater number of sites (29002 peaks) than P2-α8 (9014 peaks, **Figure 3A, Supplemental Table S4**). Consistent with this, Diffbind analysis identified 18270 P1-specific peaks, whereas most sites bound by P2-α8 were also bound by P1-α2 (**Supplemental Table S4**). HOMER motif analysis revealed that both P1 and P2 binding sites were enriched for HNF4α, CEBP, FOXA, TEAD, and KLF motifs with HNF4α motifs present in ∼50% of all binding sites (**Figure 3B**). CEBPα/β regulates hepatic differentiation^32^ while KLF4/5 regulates intestinal differentiation^33,34^, two HNF4α-dependent processes. Annotation of genomic regions bound by P1-α2 or P2-α8 revealed similar occupancy ratios between features (**Figure 3C**). Integration of our ChIPseq and RNAseq data revealed that both isoforms had greater binding at upregulated DEGs than downregulated DEGs, further supporting the observation that HNF4α serves primarily as a transcriptional activator (**Figure 3D**). We observed that P1 isoforms are stronger transcriptional repressors than P2 isoforms, as P1 occupied almost twice the amount of downregulated DEGs. Lastly, we examined binding at target genes upregulated by both isoforms from cluster 1: *VIL1* and *CLRN3* (**Figure 2C**). VIL1 is an actin-modifying protein highly expressed in epithelial cells of the intestines. CLRN3 is a key marker of the classical PDAC subtype^4^ and has biased expression in the liver and intestines. We observed greater binding of P1-α2 to the promoter and intragenic regions of *VIL1* and *CLRN3* compared to P2-α8 (**Figure 3E**), which corresponded to higher expression (**Figure 3F**).

**Figure 3:**
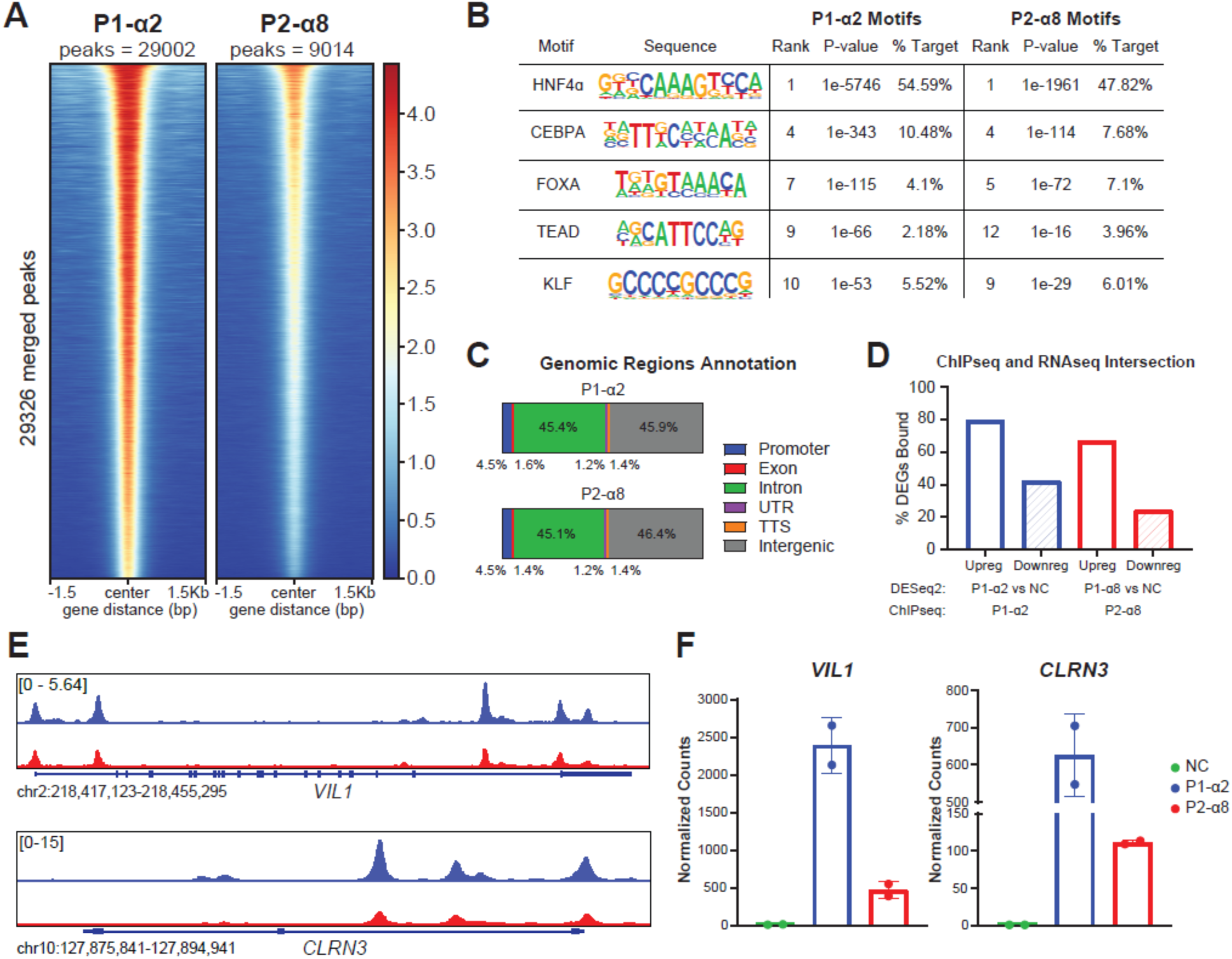
Exogenous P1-HNF4α bind at de novo loci and have increased binding at common target genes in BxPC3. HNF4α ChIP-seq analysis following expression of P1-α2 (blue) or P2-α8 (red) in BxPC3. **A.** Heatmap showing occupancy of HNF4α at merged peak regions (significant peaks called by Macs2 in at least 1 condition). Replicates were combined for each isoform (n=2). Peaks ordered by descending mean signal across all datasets. **B.** HOMER motif enrichment analysis for HNF4α, CEBPA, FOXA, TEAD, and KLF motifs. **C.** Annotation of genomic regions bound by each isoform. **D.** Percentage of significant DEGs (|log2FC|>.585, Padj<.05) bound by each isoform. **E.** ChIP-seq tracks of *VIL1 and CLRN3* (combined replicates shown, n=2). **F.** Normalized RNAseq counts of *VIL1* and *CLRN3*.

To assess the generality of these observations, we performed an integrated RNAseq/ChIP-seq analysis in Panc10.05 cells. In Panc10.05, expression of P1-α2 led to a greater number of significant DEGs (939 upregulated, 1109 downregulated) than P2-a8 (673 upregulated, 240 downregulated) when compared to control (Padj<.05, |log2FC|>1, **Supplemental Table S2**). Both isoforms induced intestinal/hepatic identities to a similar extent in Panc10.05, whereas P1 was a stronger activator of these identities in BxPC3. This may be due to cell-line-specific differences and/or higher overall levels of HNF4α in Panc10.05 (**Figure 1C**). Nevertheless, P1 inhibited the squamous/basal-like signatures to a greater extent than P2 (**Supplemental Figure S2A**). When plotting genes significantly regulated by both isoforms (Padj<.05, n=5013, **Supplemental Figure S2B**) and clustering DEGs (**Supplemental Figure S2C**), we found that P1 isoforms were stronger transcriptional regulators of common and unique target genes. Finally, Enrichr analysis revealed that genes upregulated by both isoforms were primarily enriched in intestinal and liver signatures, while genes downregulated by both were mainly enriched in non-GI tissue signatures (**Supplemental Figure S2D**).

In contrast to BxPC3, ChIP-seq analysis revealed P2-α8 bound more sites than P1-α2 in Panc10.05 (29,556 vs 16,316 peaks). (**Supplemental Figure S3A, Supplemental Table S4**). Both isoforms were enriched for similar motifs (**Supplemental Figure S3B**), types of genomic regions bound (**Supplemental Figure S3C**), and percentage of DEGs bound (**Supplemental Figure S3D**). Despite P2-α8 binding a greater number of sites, closer examination of HNF4α targets *VIL1* and *CLRN3* revealed that P1 isoforms had stronger binding (**Supplemental Figure S3E**), which corresponded with higher expression (**Supplemental Figure S3F**). Similarly, Diffbind revealed fewer differentially bound peaks than called peaks (n=4379, **Supplemental Table S4**). Altogether, these data demonstrate that exogenous P1-α2 is a stronger transcriptional regulator than P2-α8.

### CRISPRi of endogenous HNF4α reveals functional redundancies between isoforms

To complement our gain-of-function experiments, we utilized CRISPRi to knockdown expression of endogenous P1 and P2 isoforms in ASPC1 and HPAFII cells by targeting catalytically inactive Cas9 (dCas9) to their respective promoters (2 guides each, one negative control guide). We achieved robust and selective knockdown of either P1 isoforms, P2 isoforms, or both isoforms (using a dual sgRNA vector, **Figure 4A**). Interestingly, P2 knockdown led to a concomitant ∼50% increase of P1 isoforms at the RNA and protein levels (**Figure 4A-B and Supplemental Figure S4A**). In contrast, knockdown of P1 isoforms had no impact on P2 isoforms (**Figure 4A and Supplemental Figure S4B-C**). To investigate whether P2-HNF4α directly inhibits P1-HNF4α, we re-expressed exogenous P2-α8 in ASPC1 sgP2D cells, which did not inhibit the expression of P1 isoforms (**Supplemental Figure 4D**). Moreover, knockdown of P2 by shRNA (54% – 61% decrease) did not increase P1 isoform levels (**Supplemental Figure 4E**). Thus, epigenetic modification of the P2 promoter, rather than changes in P2 protein abundance, leads to increased expression of P1 isoforms.

**Figure 4.**
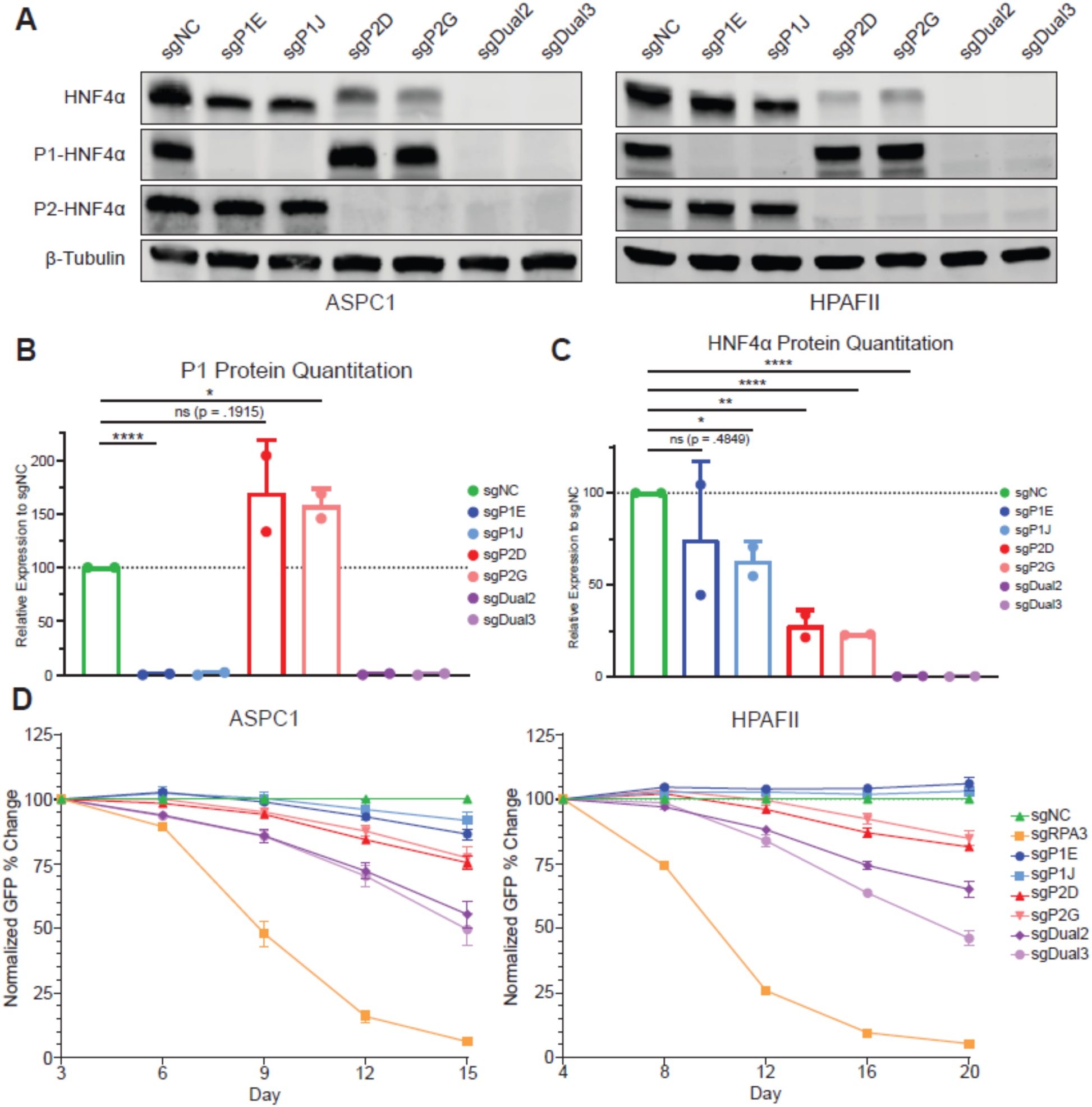
Endogenous HNF4α isoforms display functional redundancies in vitro. **A.** Representative immunoblot of HNF4α, P1-HNF4α, P2-HNF4α, or β-Tubulin after CRISPRi of P1 isoforms (sgP1), P2 isoforms (sgP2), or both P1 and P2 isoforms (sgDual) in ASPC1 and HPAFII (human PDAC cell lines), sgNC is a negative control. **B.** Quantitation of P1 protein expression in ASPC1 and HPAFII. Signal intensity was normalized to β-Tubulin then to the sgNC of each cell line, Student’s T-test, ns=not significant, *=p<.05, ****=p<.0001. **C.** Quantitation of HNF4α protein expression in ASPC1 and HPAFII. Signal intensity was normalized to β-Tubulin then to the sgNC of each cell line, Student’s T-test, ns=not significant, *=p<.05, **=p<.01, ****=p<.0001. **D.** Percent change in GFP positivity in ASPC1 and HPAFII CRISPRi competition assays after knockdown of P1, P2, or both isoforms. 3 independent experiments per cell line, percent GFP positivity was normalized to positivity at the first timepoint, then to the sgNC, sgRPA3 is a positive control, and sgNC is a negative control.

We then quantified HNF4α expression after knockdown of P1 or P2 isoforms to determine the ratio of endogenous isoforms in each cell line (**Figure 4C**). P1 isoform knockdown reduced overall HNF4α levels by 25-33%, whereas P2 isoform knockdown reduced HNF4α levels by 75% (despite the concomitant increase in P1 isoforms). Therefore, P1 and P2 proteins are expressed at a 1:3 to 1:4 ratio in these two representative PDAC cell lines.

Next, we performed CRISPRi competition assays^35^ to determine the effects of P1 and P2 knockdown on overall fitness. We transduced ASPC1 or HPAFII cells expressing dCas9 with sgRNAs also expressing GFP such that cells were ∼50% GFP positive at the first passage by flow cytometry and then tracked GFP positivity over four additional passages (**Figure 4D**). In both cell lines, knockdown of P1 isoforms led to a slight reduction in GFP positivity (∼0-10%), while knockdown of P2 isoforms led to a ∼20% reduction. In contrast, knockdown of both isoforms led to a greater decrease in GFP positivity (∼40-50%). We did not observe changes in positivity with sgNC, while sgRPA3 (positive control) led to almost a complete dropout. From these data, we conclude that a P2-high state (sgP1) is more compatible with in vitro growth than a P1-high state (sgP2), consistent with the results of expression of exogenous isoforms in HNF4α-negative cell lines (**Figure 1D**). P1 and P2 isoforms likely have partial redundancies since dual knockdown had a more severe phenotype than single knockdown of either isoform. Finally, these data show that the role of HNF4α in regulating growth is context-dependent, as knockdown of endogenous HNF4α isoforms led to a reduction in fitness, at least in the short term. In contrast, HNF4α loss at tumor initiation enhances long-term growth^12^, and exogenous expression of P1 and P2 isoforms in HNF4α-negative cell lines restrained growth^12^ (**Figure 1D**).

### Endogenous P1 isoforms are stronger transcriptional regulators than P2 isoforms

To characterize the role of endogenous HNF4α isoforms in transcription, we performed RNAseq after CRISPRi of P1, P2, or both isoforms in ASPC1 (**Figure 5**). DESeq2 analysis identified 106 DEGs in sgP1 vs. sgNC cells, 414 DEGs in sgP2 vs. sgNC cells, and 2418 DEGs in sgDual vs. sgNC cells (Padj<.05, |log2FC|>.585, **Supplemental Table S5**). The stark contrast in the number of DEGs between the knockdown of a single set of isoforms vs. both sets indicates that P1 and P2 can each partially compensate for the loss of the other isoform. GSEA of intestinal signatures did not reveal significant changes in enrichment with the loss of a single set of isoforms (**Figure 5A**). As previously reported, loss of total HNF4α led to a positive enrichment for goblet cell signatures, which we also observed in our analysis^31^. Interestingly, although not reaching statistical significance, we only observed enrichment for goblet signatures with CRISPRi of P1 and not P2, indicating that P1 isoforms may play a greater role in maintaining enterocyte and inhibiting goblet cell identity.

**Figure 5.**
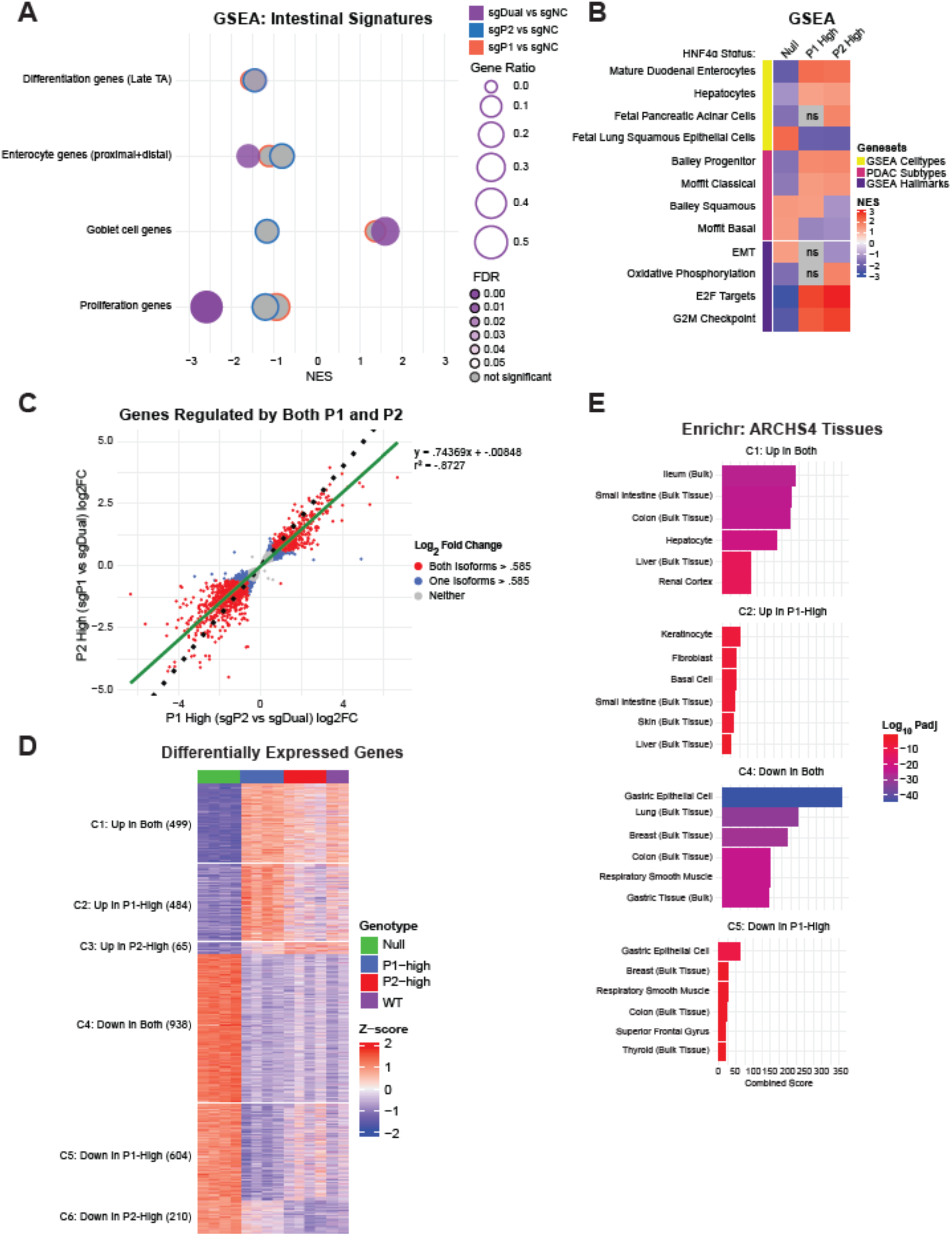
Endogenous P1 isoforms are stronger transcriptional regulators than P2 isoforms in ASPC1. RNAseq analysis following CRISPRi of P1, P2, or both isoforms in ASPC1. **A**. Dotplot of GSEA results for intestinal signatures (converted from mouse to human genes) used in Chen et al. 2019. B. Heatmap of GSEA results for the hallmark, cell type, and PDAC subtype gene sets in the following HNF4α statuses: Null (sgDual vs. sgNC), P1-high (sgP2 vs. sgDual), or P2-high (sgP1 vs. sgDual). GSEA results were significant in at least one contrast (FDR<.05), ns=not significant. **C**. Scatterplot of the log_2_ fold change of DEGs significantly regulated by both the P1-high state (sgP2 vs. sgDual) and the P2-high state (sgP1 vs. sgDual; Padj<.05). Green-line represent a linear regression model of the data (y=.74369x+-.008478, r^2^=.8727), dotted-line represents a slope of 1. **D**. Heatmap of manually clustered DEGs significantly regulated by at least one isoform in the P1-high state (sgP2 vs. sgDual) or P2-high state (sgP1 vs. sgDual; |log2FC|>1, padj<.05). **E**. Enrichr analysis results for the indicated clusters in the ARCHS4 Tissues gene set.

We then compared the transcriptome of P2-high (sgP1) and P1-high (sgP2) cells with HNF4α-null (sgDual), which identified 1712 DEGs in P2-high cells and 2526 DEGs in P1-high cells, consistent with P1 isoforms being stronger transcriptional regulators (Padj<.05, |log2FC|>1, **Supplemental Table 5**). Next, we performed GSEA on sgDual vs. sgNC, P1-high, and P2-high states (**Figure 5B**) and observed that both P1-high and P2-high were enriched for intestinal/hepatic signatures and the classical/progenitor PDAC subtypes, while loss of HNF4α led to a negative enrichment for these signatures. Dual knockdown resulted in a negative enrichment for proliferation hallmark gene sets, E2F and G2M targets, in concordance with our CRISPRi competition results. Loss of HNF4α also led to enrichment for the basal/squamous PDAC subtype and the EMT hallmark gene set. In the sgDual vs. sgNC contrast, we observed an upregulation of *SIX1* (log2FC=.73) and *SIX4* (log2FC=1.28, **Supplemental Table 5**), lineage specifiers that regulate mesodermal differentiation and the basal-like subtype as previously reported in our GEMM^12^.

We then plotted DEGs regulated by both the P1-high and P2-high states (n=4927, **Figure 5C**) and fit a linear model of the data, which had a slope of .74, indicating that for a given common target gene, P1 is a stronger transcriptional regulator. Next, we manually clustered and plotted a heatmap of all significant DEGs from the P1-high and P2-high states based on the directionality of gene expression (Padj<.05, |log2FC|>1), resulting in 6 distinct clusters (**Figure 5D, Supplemental Table S6**). P1 isoforms regulated a much greater number of unique DEGs (C2 and C5) than P2 isoforms (C3 and C6). Of note, in cluster 2 (up in P1-high), the P1-high state has a higher z-score than in the WT (both isoforms present), which may be the result of the compensatory increase in absolute P1 isoforms and/or reflect an ability of P2 to dampen P1 activity via heterodimerization^20^ (**Figure 4A-B**). Enrichr analysis for the ARCHS4 Tissues gene sets (**Figure 5E, Supplemental Table S6**) revealed that C1 (up in both isoforms) was enriched for canonical intestinal and liver cell types, while C4 (repressed by both isoforms) was enriched for non-GI cell types, including bulk breast and lung signatures. While we also observed some enrichment for gastric and bulk colon signatures in C4, closer investigation revealed that the downregulated genes driving this enrichment are broadly expressed in multiple tissue types. DEGs associated with the P1-high state (C2 and C5) exhibited minimal tissue specificity (low overall score, absence of coherent pattern of tissue/cell type enrichment).

To compare genome occupancy of endogenous HNF4α isoforms, we performed HNF4α ChIP-seq in ASPC1 (**Figure 6**) in sgNC (both isoforms), sgP1E (P2-high), and sgP2D (P1-high) using a pan-HNF4α antibody. ChIP-seq revealed that the P1-high (10,305 peaks) and P2-high states (9614 peaks) had a similar number of HNF4α binding sites despite lower levels of HNF4α protein in P1-high cells (**Figure 4A-B**). Both had fewer peaks than the control, as expected, given higher overall HNF4α levels in control cells (13,743 peaks, **Figure 6A, Supplemental Table S7**). Diffbind analysis between the P1-high and P2-high states exhibited very few differentially bound sites (**Supplemental Table S7**). HOMER motif analysis revealed similar enrichment for the HNF4α motif, but the control and P1-high state had greater enrichment for the KLF and TEAD motif than the P2-high state (**Figure 6B**). Annotation of the different genomic regions bound by control, P1-high or P2-high, indicated similar binding ratios to genomic features (**Figure 6C**). Integration of our endogenous RNAseq and ChIPseq revealed that HNF4α isoforms primarily serve as transcriptional activators as there is a greater percentage of genes bound by HNF4α isoforms that are downregulated after CRISPRi (**Figure 6D**). Lastly, we examined HNF4α binding (**Figure 6E**) and changes in RNA expression levels (**Figure 6F**) of *VIL1* and *CLRN3*. We found that both the P1-high and P2-high states had comparable levels of HNF4α binding at the promoter and intragenic regions of these targets. CRISPRi of either P1 or P2 isoforms resulted in a similar decrease in expression of *VIL1* but not *CLRN3*. However, the loss of both isoforms (sgDual) led to a further reduction in the expression levels of both genes.

**Figure 6.**
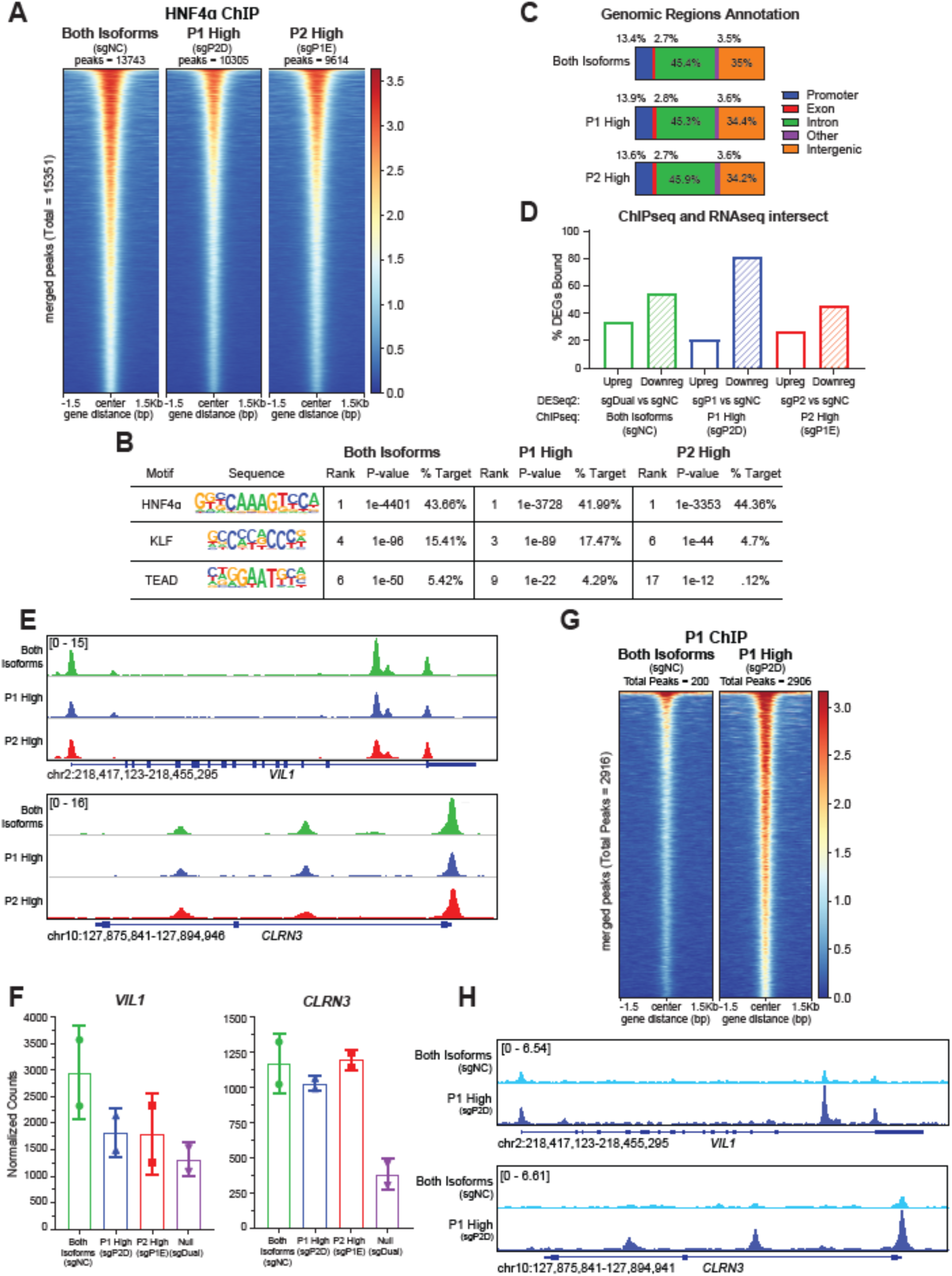
HNF4α isoforms exhibit compensatory binding following the loss of one set of isoforms in ASPC1. HNF4α ChIP-seq analysis in ASPC1 cells expressing both isoforms (sgNC, green), P1-high (sgP2D, blue), or P2-high (sgP1E, red). **A.** Heatmap showing occupancy of HNF4α at merged peak regions (significant peaks called by Macs2 in at least 1 condition). Replicates were combined for each isoform (n=2). Peaks ordered by descending mean signal across all datasets. **B.** HOMER motif enrichment analysis for HNF4α, KLF, and TEAD motifs. **C.** Annotation of genomic regions bound by each isoform. **D.** Percentage of significant DEGs (|log2FC|>.585, Padj<.05) bound by each isoform. **E.** HNF4α ChIP-seq tracks of *VIL1 and CLRN3* (combined replicates shown, n=2). **F.** Normalized RNAseq counts of *VIL1* and *CLRN3*. **G.** Heatmap showing occupancy of P1 isoforms at merged peak regions (significant peaks called by Macs2 in at least 1 condition). Replicates were combined for each isoform (n=2). Peaks ordered by descending mean signal across all datasets. **H.** P1 ChIP-seq tracks of *VIL1 and CLRN3* (combined replicates shown, n=2).

To further characterize chromatin binding patterns of endogenous P1-HNF4α and P2-HNF4α, we sought to perform ChIP-seq with the isoform-specific P1 and P2 antibodies. We first performed ChIP-qPCR in BxPC3 cells expressing P1-α2 or P2-α8 to validate the antibodies (**Supplemental Figure S5A**). With the P1 antibody, we were able to observe enrichment over input at *HNF1A* and *VIL1* in P1-α2 cells but not in the negative control region, NC2, or in P2-α8 cells indicating the P1 antibody was specific and suitable for ChIP-seq. We did not observe any differences in enrichment over input with the P2 antibody (**Supplemental Figure S5A**). ChIP-qPCR further confirmed the sensitivity and specificity of the P1 antibody (**Supplemental Figure S5B**) and the lack thereof with the P2 antibody in ASPC1 (**Supplemental Figure S5C**). While P1-ChIP in ASPC1 sgNC revealed a small number of peaks (200 peaks), P1-ChIP in P1-high cells revealed a robust number of peaks (2906 peaks) (**Figure 6G**). Closer examination of HNF4α target genes, *VIL1* and *CLRN3,* revealed an increase in P1 binding at the promoter and intragenic regions in P1-high compared to the WT (**Figure 6H**). These data further support our finding that P1 isoform levels and activity increase after CRISPRi of P2. In summary, while P1-HNF4α and P2-HNF4α can compensate for the loss of their reciprocal set of isoforms, P1-isoforms are stronger transcriptional regulators since a lower amount of P1 isoforms is sufficient to compensate for the loss of a greater amount of P2 isoforms.

### Integrated RNAseq and ChIP-seq analysis reveals core HNF4α target genes

We integrated RNAseq datasets from BxPC3, Panc10.05, and ASPC1 to identify a list of consensus HNF4α-regulated genes in PDAC (**Supplemental Figure S6, Supplemental Table S8**). There were 82 genes upregulated by P1 and 36 genes upregulated by P2 in all three datasets (**Supplemental Figure S6A-B**). There were 25 genes downregulated by P1 and 2 genes downregulated by P2 in all three datasets (**Supplemental Figure S6C-D**). When contrasting the consensus P1– and P2-regulated genes, we observed that almost all P2-regulated genes were also regulated by P1 (**Supplemental Figure S6E**). Enrichr analysis of the genes upregulated by both isoforms (**Supplemental Figure S6F, Supplemental Table S8**) revealed a robust enrichment for intestinal/hepatic signature as evidenced by a combined score 3-fold higher than our BxPC3 Enrichr analysis (**Figure 2E**). Like our previous observations, Enrichr analysis of genes unique to P1 isoforms resulted in a slight but significant enrichment for intestinal/hepatic signatures.

We then examined DNA binding and gene expression for each category of HNF4α target genes to better understand how HNF4α affects transcription. We plotted ChIP-seq peaks and normalized counts of a representative P1-upregulated gene: *CES2* (**Supplemental Figure S7A-B**), P1/P2-upregulated gene: *EPS8L3* (**Supplemental Figure S7C-D**), P2-upregulated gene: *AZGP1* (**Supplemental Figure S7E-F**), and P1/P2-downregulated gene: *IFI27* (**Supplemental Figure S7G-H**). We observed shared HNF4α peaks in all cell lines in all three categories of upregulated target genes. Upon closer scrutiny, *AZGP1* was identified as a P2-upregulated gene in the Panc10.05 RNAseq data only (**Supplemental Figure S7F**). However, there was still greater binding of P1 at the promoter, indicating that the stronger upregulation of *AZGP1* may be due to non-specific effects of P2 expression (**Supplemental Figure S7F**). Although little HNF4α binding was observed at *IFI27,* a gene downregulated by both isoforms (**Supplemental Figure S7G**), the expression of this gene still trended lower in P1-high than P2-high states (**Supplemental Figure S7H**). This pattern was also observed at other downregulated genes (data not shown), indicating that P1 and P2 isoforms do not predominantly inhibit gene expression by direct DNA binding.

### CRISPRi of HNF4α isoforms is insufficient to alter enhancer-promoter contacts at HNF4α target genes

It has been previously reported that HNF4α/γ are required for chromatin looping in intestinal villi^36^. Based on the spatial distribution of HNF4α isoforms in the intestinal epithelium (P1 – differentiated colonocytes; P2 – proliferating cells)^26,27^, we investigated whether the increased transcriptional activity of P1 isoforms is due to its ability to regulate enhancer-promoter (E-P) interactions by performing H3K27ac Hi-ChIP in ASPC1 cells. Globally, we did not observe significant changes in E-P contacts at HNF4α target genes in either the single or dual knockdowns (**Supplemental Figure S8A**). Investigation of target genes bound by HNF4α also did not yield significant changes in E-P contacts (**Supplemental Figure S8B**). CRISPRi successfully disrupted E-P interactions at the *HNF4A* locus at the respective promoters (**Supplemental Figure S8C**), supporting the sensitivity of the HiChIP assays. However, the expression increase in P1 isoforms after knockdown of P2 was not due to the rewiring of chromatin interactions as E-P contacts remain unchanged between sgP2D and sgNC (**Supplemental Figure S8C**). The expression increase in P1 isoforms may be due to released transcriptional interference from P2 isoforms after CRISPRi of P2. Lastly, a closer examination of the *CLRN3* locus, a canonical HNF4α target gene, did not reveal significant changes in E-P contacts with knockdown of a single set of isoforms but revealed a significant decrease with knockdown of total HNF4α (**Supplemental Figure S8D-E**). These data suggest that P1 and P2 isoforms do not differentially regulate E-P contacts, at least in this representative cell line, and that the difference in transcriptional activity may be downstream of chromatin alterations (e.g., differential P1/P2 binding and recruitment of cofactors).

## DISCUSSION

While previous studies have established the importance of HNF4α as a key regulator of the classical PDAC subtype, the functional roles of its P1– and P2-HNF4α isoforms remained unclear. In this study, we characterized P1 and P2 expression in primary PDAC and IPMNs and investigated their effects on PDAC cell growth and gene expression. Utilizing complementary gain-of-function and CRISPRi approaches, we demonstrated that P1-HNF4α isoforms are less compatible with growth and are stronger transcriptional regulators than P2-HNF4α.

In both exogenous and endogenous HNF4α models, we found that P1 isoforms are stronger transcriptional regulators compared to P2 isoforms. Although we initially hypothesized that P1 and P2 isoforms may drive distinct identities (i.e., gastric vs. intestinal) based on their expression in normal tissue, our study revealed that both isoforms induce intestinal/hepatic identity programs, with P1 acting as a stronger lineage specifier than P2. This difference in transcriptional activity is likely attributable to the structure of the P1 and P2 first exons. The P1 exon 1 contains an AF-1 transactivation domain, which may enhance transcriptional activity through interactions with cofactors. Supporting this, a recent study expressed all 12 isoforms of HNF4α in a human colon cancer cell line and found that P1 isoforms were more transcriptionally active than P2 isoforms^21^. Downstream analysis of proteins that interact with HNF4α revealed both shared and isoform-specific cofactors^21^. These findings suggest that an unbiased protein interaction assay in PDAC may reveal cofactor interactions that explain the increased transcriptional activity of P1 isoforms.

In both exogenous and endogenous models, we demonstrated that P1 isoforms are less compatible with growth than P2 isoforms. However, the role of HNF4α in regulating PDAC growth appears to be context-specific. While exogenous HNF4α restrained growth, knockdown of endogenous HNF4α did not increase growth, indicating a nuanced role in classical PDAC cell lines. One explanation is that PDAC cells expressing endogenous HNF4α have adapted to its presence and rely on HNF4α-regulated cellular processes. Removing HNF4α in this context disrupts homeostasis, resulting in a loss of fitness. These results complement our previous studies of HNF4α loss at tumor initiation, where deletion at the start of tumor progression increased tumor burden in a GEMM^12^. Developing tumor cells, independent from HNF4α’s effects in maintaining differentiation and metabolism, adopt a more proliferative alternative state in its absence. A similar context-dependent role in regulating growth has been described for PDX1, a master regulator of pancreatic embryogenesis and β-cells. PDX1 plays an oncogenic role in the progression to PanINs and PDAC but is lost in a subset of PDAC tumors that then transition to a squamous subtype^37^.

Our results raise two critical questions: Why are P1 isoforms less compatible with growth than P2 isoforms in PDAC, and since P1 isoforms are less compatible with growth, why are they expressed in PDAC? Although we did not identify P1-specific target gene(s) that represses growth in PDAC, the data suggest two possible mechanisms. P1 isoforms may be less compatible with growth than P2 isoforms through stronger transcriptional regulation of shared target genes or regulation of a unique set of genes. These mechanisms are not mutually exclusive and may both contribute to some degree. This regulation may happen directly via stronger DNA binding by P1 isoforms or indirectly through activation of downstream transcription factors. We did not observe robust DNA binding at downregulated target genes, suggesting that HNF4α may not act as a direct transcriptional repressor. Despite this, we found that exogenous P1 isoforms were more negatively enriched for proliferative intestinal cell signatures than P2 isoforms. Ultimately, P1 isoforms may be less compatible with growth through the transcriptional regulation of multiple genes, which in their totality attenuates growth.

There are two potential explanations as to why PDAC cells express P1 isoforms despite it being less compatible with growth. First, P1 isoforms may originate from IPMNs that progressed to PDAC, as documented by our study and a recently published study^29^. We speculate that P1 isoforms may play a growth-promoting role in the development of dysplasia/IPMNs but are no longer needed in cancer. Similar observations have been made in esophageal adenocarcinoma and gastric cancer, in which CDX2 expression is upregulated in Barrett’s esophagus/intestinal metaplasia but lost at subsequent steps in progression^38^. However, it is unlikely that IPMNs are the only source of P1 expression in PDAC since they are not the predominant precursor lesion to PDAC, and most PDAC cases we evaluated were not associated with IPMNs. It is also possible that P1 expression may stochastically arise in PDAC tumors because of transcriptional or epigenetic events during tumor progression. In this second scenario, P1 expression may arise as a passenger event where drivers of PDAC progression inadvertently upregulate P1 isoforms. Further experiments are needed to elucidate the exact mechanisms accounting for the heterogeneity of P1 expression in primary human PDAC. Evidence in other malignancies suggests post-translational modifications may be important in regulating the expression of P1 isoforms. In colorectal carcinoma, SRC has been demonstrated to directly phosphorylate exon 1 of P1, leading to its degradation^39^, and β-catenin has been shown to repress the expression of P1 isoforms^27^. Although SRC and β-Catenin are not the primary oncogenic drivers of PDAC, post-translational mechanisms may account for the cell-to-cell variability in P1 expression in our IHC analysis.

While HNF4α isoforms share some functional redundancies in PDAC, P1 and P2 isoforms may display more pronounced differential activity in other gastrointestinal malignancies. For instance, in gastric cancer, P1 isoforms are correlated with CDX2 expression and intestinal metaplasia^40^, whereas no such correlation was observed in intestinal-type IPMNs. Additional studies in gastrointestinal tissues that undergo intestinal metaplasia during tumor progression may uncover additional insights into the differential activity of P1– and P2-HNF4α. Notably, the CRISPRi reagents generated in this study are a significant improvement over RNAi-based approaches, offering a valuable tool for further investigations into HNF4α isoform biology. In summary, our study provides a comprehensive analysis of the differential roles of P1 and P2 isoforms in PDAC, and our data show that careful consideration of gene isoforms is necessary when studying endodermal lineage specifiers, as they may have distinct roles in tumor biology.

## MATERIALS AND METHODS

### Primary human tumors

PDAC and IPMN whole sections from FFPE blocks, and associated clinical/pathological data, were obtained via the HCI Biorepository under IRB protocols approved by the University of Utah. Slides of human material were reviewed by two board-certified pathologists (K.A. and E.L.S.).

Additional materials and methods are provided in supplemental information.

## Acknowledgments

We are grateful to members of the Snyder lab for helpful suggestions and comments. We thank Charles Murtaugh for providing ASPC1 and HPAFII cell lines, Xiaoyang Zhang for providing Lenti-MeCP2-Blasticidin plasmid and for CRISPRi and HiChIP expertise, Olaf Klingbeil for dual sgRNA cloning expertise, and Luke Torre-Healy for analysis of HNF4α isoform expression in PDAC cell lines. The schematic for the *HNF4A* loci for this publication was created with Biorender.com.

## Contributors

PF and ELS designed experiments. PF, AK, and XYZ performed experiments. PF, ERW, CS, and ELS analyzed data. ELS and KA performed histopathological review. PF and ELS wrote the manuscript. All authors discussed the results and reviewed and revised the manuscript.

## RESOURCE AVAILABILITY

### Lead contact

Further information and requests for resources and reagents should be directed to and will be fulfilled by the lead contact, Eric Snyder

### Materials availability

Plasmids and cell lines generated in this study are available upon request.

### Data and code availability

- Bulk RNA-seq and ChIP-seq data have been deposited at Gene Expression Omnibus GSE284709 and are available as of the date of publication.
- This manuscript does not report original code.
- Any additional information required to reanalyze the data reported in this manuscript is available from the Lead Contact upon request.

