## Supplemental figures and methods for "Differential control of growth and identity by HNF4α isoforms in pancreatic ductal adenocarcinoma"

**MATERIALS AND METHODS**

**Cell lines**

BxPC3, Panc10.05, ASPC1, HPAFII, and HEK-293T were cultured in RPMI-1640 supplemented with 10% FBS (VWR), 1% Glutamax (Gibco), and 2.5ug/mL Plasmocin (to maintain cell cultures free of mycoplasma; InvivoGen). All cell lines were tested periodically for mycoplasma contamination. The identity of cell lines was verified by STR genotyping through the University of Utah DNA Sequencing Core.

**Plasmids and lentiviral production**

Dox-inducible, pCW22 HNF4α2 (P1) and HNF4α8 (P2) were previously published in Camolotto et al. Gut 2020^1^. HNF4α2 (P1) and HNF4α8 (P2) were cloned into a dox-inducible, pCW57 Blast lentiviral expression vector (addgene #80921) via AgeI and BamHI restriction enzyme cloning. Lenti-Mecp2-Blast-dCas9 was a gift from X.Y.Z.^2^. CRISPRi guide RNAs were designed to target + and – 300 bp from the TSS of P2 and P1 promoters and cloned into LRG2.1 Puro (addgene #125594) lentiviral expression vector via BsmBI restriction enzyme cloning. Dual guide RNA expression vectors were cloned by Gibson Assembly (NEB) by insertion of a gBlock gene fragment (IDT; homology arm – guide #1 – guide backbone – alternative species U6 promoter – guide #2 – homology arm) into the LRG2.1 Puro backbone. CRISPRi negative control guide was cloned from previously published negative control sequence^2^. shP2 was cloned into pLKO.1 – TRC cloning vector (addgene #10878) from previously published shRNA sequence^3^. All CRISPRi guide RNA and shRNA sequences are listed in **Supplemental Table S9**.

Lentiviruses were produced by transfection of HEK-293T cells with TransIT-293 (Mirus Bio) transfection reagent, lentiviral-plasmids, packaging (Δ8.9 (gag/pol)) and envelope vectors (VSV-G), and Opti-MEM (Gibco) for 24-hours. The supernatant was collected at 48- and 72-hours post-transfection and filtered using 0.45 μM filters.

**RNA extraction, cDNA synthesis, and qPCR**

RNA was isolated via Trizol-chloroform extraction followed by column-based purification. The aqueous phase was brought to a final concentration of 35% ethanol, and RNA was purified using the PureLink RNA Mini Kit (ThermoFisher Scientific) according to the manufacturer’s specifications.

cDNA was synthesized from Trizol-extracted RNA using LunaScript RT SuperMix (NEB, M3010) according to the manufacturer’s specifications. qPCR was performed on cDNA using Luna Universal qPCR Master Mix (NEB, M3003) according to the manufacturer’s specifications; 35 cycles were used for the denaturation and extension steps. Transcript levels were normalized to PPIA and quantitated by the ΔΔCt method. All SYBR primers used are listed in **Supplemental Table S9**.

**Immunoblotting**

Protein was extracted by lysing cells on ice for 20 mins in RIPA buffer (50 mM Tris HCl pH 7.4, 150 mM NaCl, 0.1% (w/v) sodium dodecyl sulfate, 0.5% (w/v) sodium deoxycholate, 1% (v/v) Triton X-100) plus Pierce Protease Inhibitor (PPI, ThermoFisher Scientific, A32959). Protein was isolated by centrifugation for 10 minutes at 4°C and collection of subsequent supernatant. Protein concentration was measured by Pierce Bradford Protein Assay Kit (ThermoFisher Scientific, 23200). 10-20 μg of protein per sample were resolved on Tris-Glycine precast gels (SMOBIO, QP4510) and transferred to a nitrocellulose membrane (ThermoFisher Scientific, 88018). Membranes were probed overnight with antibodies against HNF4α (1:500, CST, C11F12), P1-HNF4α, (1:250, R&D, PP-K9218-00), HNF4α-P2 (1:250, R&D, PP-H6939-00), β-Tubulin (1:1000, DSHB, E7), and Vinculin (1:20,000, Abcam, 129002). Membranes were subsequently probed with fluorescent-conjugated goat-anti-mouse-800-CW and goat-anti-rabbit-680-RD secondary antibodies (1:20,000, LI-COR Biosciences) for 1 hour. Protein detection was performed using Odyssey CLx Imaging System (LI-COR Biosciences).

**Incucyte**

Cell growth was monitored by changes in confluence over 120 hours via a live cell imaging system (IncuCyte). BxPC3 and Panc10.05 cells were treated with doxycycline (Sigma-Aldrich, D9891) at the following concentrations (BxPC3 α2/α8/empty vector – 500ng/mL; Panc10.05 α2/gfp – 25ng/mL; α8 – 12.5ng/mL for 72 hours prior to seeding. 4,000 BxPC3 cells and 6,000 Panc10.05 cells were seeded per well (8 wells per condition). Doxycycline was replenished at 48- and 96-hours. Results were normalized to the initial seeding density at 24 hours.

**CRISPRi competition**

ASPC1 and HPAFII were first transduced with dCas9. Guide RNA viruses were titered such that cells were ~40-60% positive at initial measurement (ASPC1 – day 3; HPAFII – day 4) by flow cytometry. Flow cytometry was performed using BD Fortessa. Percent GFP positive were then monitored at four additional time points (ASPC1 – every 3 days; HPAFII – every 4 days). 10,000 events per condition at each time point were recorded for ASPC1, and 5,000 events per condition at each time point were recorded for HPAFII. Results were first normalized to the initial % GFP positive of each sample and then to the negative control at each time point.

**Immunohistochemistry**

Immunohistochemistry (IHC) was performed manually on Sequenza slide staining racks (Thermo Fisher Scientific). Sections were treated with Bloxall (peroxidase block; Vector Labs) followed by horse serum (protein block; Vector Labs), primary antibody, and HRP-polymer-conjugated secondary antibody (anti-Rabbit). The slides were developed with Impact DAB (Vector Labs) and counterstained with hematoxylin. Slides were stained with antibodies to HNF4α (1:500, CST, C11F12) and CDX2 (1:500, CST, D11D10). HNF4α-P1 (1:250, R&D, PP-K9218-00) and HNF4α-P2 (1:500, R&D, PP-H6939-00) IHC was performed using ImmPRESS Excel Amplified Polymer Staining Kit, Anti-Mouse IgG, Peroxidase (MP-7602) in Sequenza slide staining racks according to manufacturer’s specifications. Images were taken on a Nikon Eclipse Ni-U microscope with a DS- Ri2 camera and NIS-Elements software. Histological analyses were performed on hematoxylin and eosin-stained, and IHC-stained slides using NIS-Elements software. All histopathologic analysis was performed by a board-certified anatomic pathologist (E.L.S.).

**RNA Sequencing**

RNA was collected in biological duplicates after 7 days of dox-treatment to induce expression of exogenous P1, P2, or control in BxPC3 and Panc10.05 or after CRISPRi knockdown of P1, P2, both isoforms, or control in ASPC1. Library preparation was performed using NEBNext Ultra II Directional RNA Library Prep with poly(A) mRNA isolation. Sequencing was performed using Illumina NovaSeq 6000 (150 x 150 bp paired-end sequencing, 25 million reads per sample).

**RNA-seq data processing and analysis**

The human hg38 genome and gene feature files were downloaded from Ensembl and a reference database was created using STAR version 2.7.6a^4^. Optical duplicates were removed from NovaSeq runs via Clumpify v38.34^5^. Reads were trimmed of adapters and aligned to the reference database using STAR in two-pass mode to output a BAM file sorted by coordinates. Mapped reads were assigned to annotated genes using featureCounts version 1.6.3^6^. Raw counts were filtered to remove features with zero counts and features with five or fewer reads in every sample. DEGs were identified using the hciR package (<https://github.com/HuntsmanCancerInstitute/hciR>) with a 5% false discovery rate and DESeq2 version 1.34.0^7^. fGSEA (hciR package) was run in BxPC3, Panc10.05, and ASPC1 with the differentially expressed gene list generated from DESeq2 and the following MSigDB gene sets: Hallmarks and C8 as well as a custom gene set of PDAC subtype signatures. Gene sets smaller than 15 and larger than 500 were excluded from analysis. Heatmaps were generated using ComplexHeatmap^8^. Significant, differentially expressed genes (log2FC > 1 or < -1; pdaj <.05) were clustered by intersecting DEGs regulated by P1 and/or P2. Pathway analysis was performed on each cluster using Enrichr^9^.

**ChIP sequencing**

For exogenous ChIP-seq, BxPC3 and Panc10.05 cells were treated with dox for 7 days to induce expression of P1 or P2. For endogenous ChIP-seq, P1 or P2 expression was knocked down by CRISPRi in ASPC1; sgNC was used to ChIP for both isoforms. All ChIP-seq experiments were performed in biological duplicates. ~10 million cells were collected per condition. Cells were washed in cold PBS and resuspended in 10 mL of DSG buffer (1X PBS, 1 mM MgCl_2_). .25M Disuccinimidyl Glutarate (DSG) was added, and cells were rotated at room temperature for 35 minutes. Methanol-free formaldehyde was then added to a final concentration of 1%, and cells were crosslinked for 10 minutes. The cross-linking reaction was quenched with the addition of glycine (to a final concentration of 125 mM) and rotation at room temperature for 5 minutes. Cells were washed with cold PBS, and the cell pellet was snap-frozen in liquid nitrogen and stored at -80°C.

Cell pellets were thawed on ice for 5 minutes and then lysed in 1 mL of Farnham lysis buffer (5 mM PIPES pH 8.0, 85 mM KCl, 0.5% NP40) with Pierce Protease Inhibitor (PPI, ThermoFisher Scientific, A32959). Samples were centrifuged at 4°C then resuspended in 1mL of RIPA lysis buffer (1X PBS, 1% NP40, 0.5% sodium deoxycholate, 0.1% sodium dodecyl sulfate) with PPI. Chromatin was sonicated with QSonica Q800R (pulse: 30s on/30s off; sonication time: 20 minutes; amplitude: 70%). After sonication, an input was collected from each sample of sheared chromatin. Chromatin from each sample was then immunoprecipitated overnight with 5ug of the following antibodies (per sample) premixed with Protein G Dynabeads (ThermoFisher Scientific, 10004D): HNF4A (R&D, PP-H1415-0C, mouse-monoclonal), P1-HNF4α (R&D, PP-K9218-00, mouse-monoclonal) or P2-HNF4α (R&D, PP-H6939-00, mouse-monoclonal).

Library preparation was performed using ChIP-seq with NEBNext DNA Ultra II library prep kit using Unique Molecular Indexes (UMIs). Sequencing was performed using Illumina NovaSeq 6000 (150 x 150 bp paired-end sequencing, 25 million reads per sample). For ChIP-qPCR assays, primers were designed targeting HNF4α binding sites identified from publicly available ChIP-seq data from ENCODE (<https://www.encodeproject.org/>) on Integrative Genomics Viewer (IGV; <https://software.broadinstitute.org/software/igv/>). All ChIP-qPCR primers are listed in **Supplemental Table S9**.

**ChIP-seq data processing and analysis**

Fastq alignments were pre-processed with the merge_umi_fastq application from the UMIScripts package (<https://github.com/HuntsmanCancerInstitute/UMIScripts>) to associate the UMI sequence, provided as a third Fastq file, into the read comment. Reads were aligned using Bowtie2 v2.2.9^10^ to the standard chromosomes of the human genome (version hg38). Duplicate alignments based on the UMI code were removed using the bam_umi_dedup application (UMIScripts) allowing for 1 mismatch. Peaks were called using MACS2 v2.2.7^11^ with a significance of q-value < 1e-8 in exogenous HNF4α ChIP and q-value < .01 in endogenous HNF4α ChIP. Coverage tracks were generated with MACS2 as Reads Per Million. Input libraries were obtained from all samples and were used as controls for each ChIP-seq experiment. All ChIP-seq experiments were performed in biological duplicates. Peaks called in both biological replicates were identified using Bedtools v2.28.0^12^ with a 1-bp minimum overlap to generate a consensus list of peaks for downstream analysis. Genomic annotation of binding sites was performed using HOMER^13^. IGV was used to visualize ChIP-seq peaks. Differential ChIP-seq peaks were identified using the Diffbind package v3.4.11 (https://bioconductor.org/packages/release/bioc/vignettes/DiffBind/inst/doc/DiffBind.pdf) with a q-value cutoff < 0.05 using DESeq2 for the analysis method. Tornado plots were generated using deeptools v3.5.1^14^.

**HiChIP**

For HiChIP, 6 million cells were collected per replicate. Cells were fixed in 1% formaldehyde for 10 minutes, then quenched with glycine to a final concentration of 125 mM. HiChIP was performed as described (<https://www.nature.com/articles/nmeth.3999>) with minor modifications as described (<https://www.nature.com/articles/s41467-021-27055-4>). Cross-linked chromatin was digested with the MboI restriction enzyme followed by end-repair with dNTPs including biotin labeled dATP, ligation using T4 DNA ligase, and sonication to obtain 1kb chromatin fragments using Qsonica (Q800). To enrich for chromatin interactions occurring at active regulatory elements, an anti-H3K27ac antibody (Abcam, ab4729, rabbit polyclonal, 7.5ug/HiChIP) was used for DNA fragment capture. Streptavidin magnetic beads were used to pull down ligated DNA fragments and HiChIP libraries were prepared using Illumina Tagment DNA Enzyme and Buffer Kit. Sequencing was performed with NovaSeq X at the High-Throughput Genomics core at the Huntsman Cancer Institute, University of Utah.

The paired-end HiChIP sequencing reads were aligned to the human genome hg38 with the HiC-Pro pipeline (<https://genomebiology.biomedcentral.com/articles/10.1186/s13059-015-0831-x>). HiC-DC+ was used to call chromatin loops by binning the genome into 5kb bins (<https://www.nature.com/articles/s41467-021-23749-x#citeas>). Chromatin loops were removed if the qvalue > 0.05 or anchors overlapped with ENCODE blacklist regions (<https://www.nature.com/articles/s41598-019-45839-z>). BEDTools pairToBed was used to identify enhancer promoter interactions, loops with only one anchor overlapping with transcription start site (TSS) of a gene (<https://academic.oup.com/bioinformatics/article/26/6/841/244688>). HiChIP data was visualized at the resolution of 5kb using the R package gTrack (<https://github.com/mskilab-org/gTrack>).

**Quantification and Statistical Analysis**

All graphing and statistical analysis were performed with PRISM software or R, with all graphs showing the mean and either the standard deviation or standard error of the mean. The statistical details can be found in the corresponding figure legend. All NGS statistical analysis was performed according to published pipeline protocols cited, with a statistical significance cutoff of padj < 0.05.

**SUPPLEMENTAL FIGURE S1**

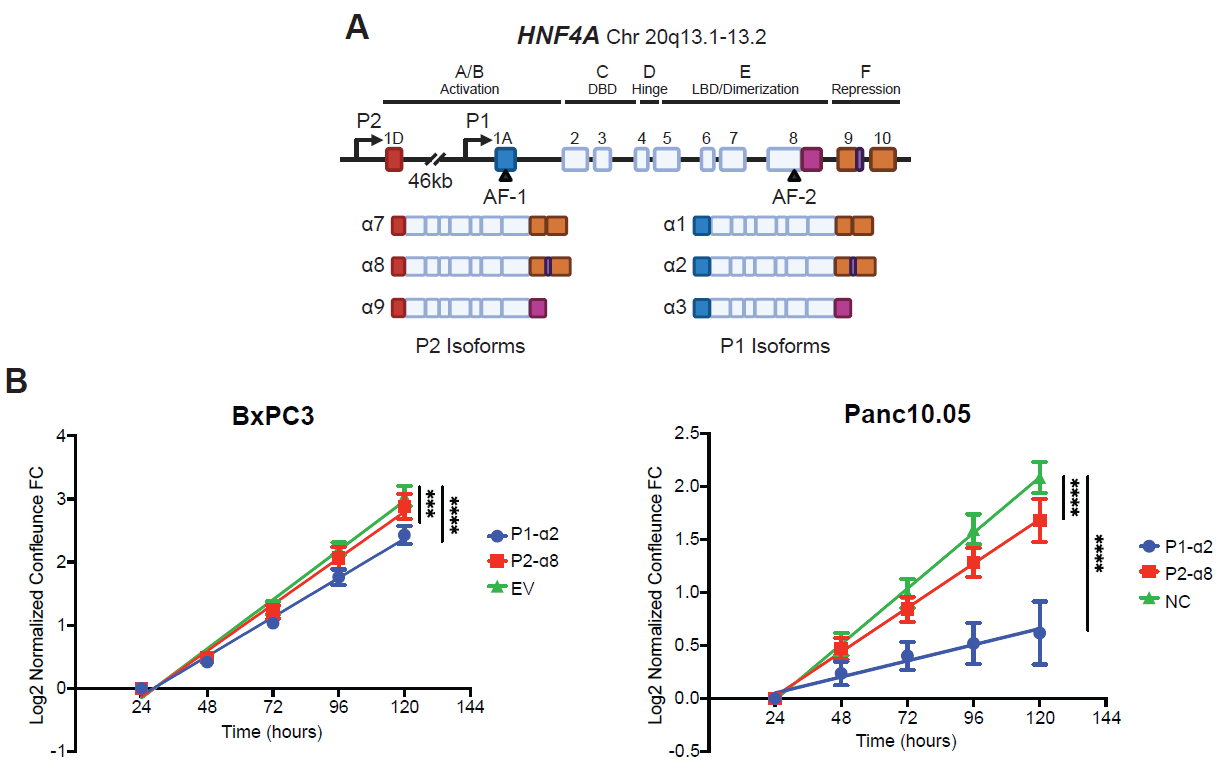

**Supplemental Figure S1: Exogenous expression of P1-HNF4α restrains tumor growth in vitro.**

**A.** Schematic of the *HNF4α* loci with canonical P1 and P2 isoforms.

**B.** Log_2_ transformation of incucyte growth assays in BxPC3 and Panc10.05 cells for ANCOVA significance of slope calculations (***=p<.001, ****=p<.0001, **Figure 1E**).

**SUPPLEMENTAL FIGURE S2**

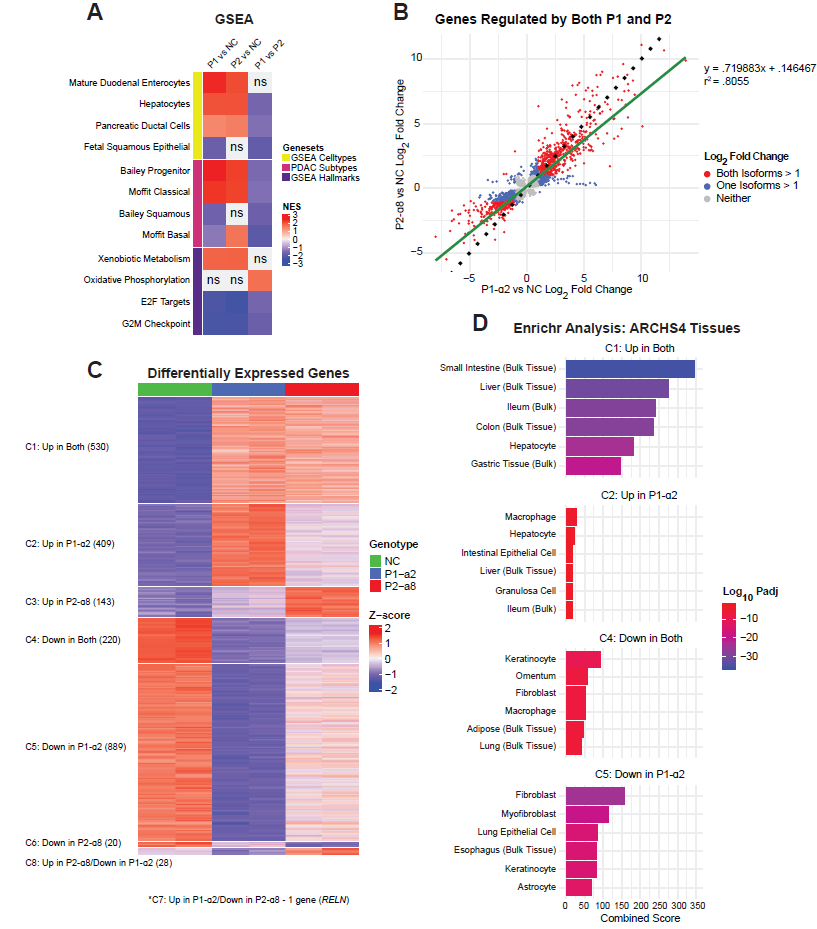

**Supplemental Figure S2: Exogenous P1-HNF4α is a stronger transcriptional regulator than P2-HNF4α in Panc10.05.**

RNAseq analysis following expression of P1-α2 or P2-α8 in Panc10.05.

**A.** Heatmap of GSEA results for the hallmark, cell type, and PDAC subtype gene sets in the following contrasts: P1-α2 vs. NC, P2-α8 vs. NC, and P1-α2 vs. P2-α8. In the P1-α2 vs P2-α8 contrast, a positive NES score indicates enrichment in P1, whereas a negative NES score indicates enrichment in P2. GSEA results were significant in at least one contrast (FDR<.05), ns=not significant.

**B.** Scatterplot of the log_2_ fold change of DEGs significantly regulated by both isoforms (padj <.05). Green-line represents a linear regression model of the data (y = .719883x + .146467, r^2^ = .8055), dotted-line represents a slope of 1.

**C.** Heatmap of manually clustered DEGs significantly regulated in at least one contrast (log2FC > 1 or < -1 and padj < .05).

**D.** Enrichr analysis results for the indicated clusters in the ARCHS4 Tissues geneset.

**SUPPLEMENTAL FIGURE S3**

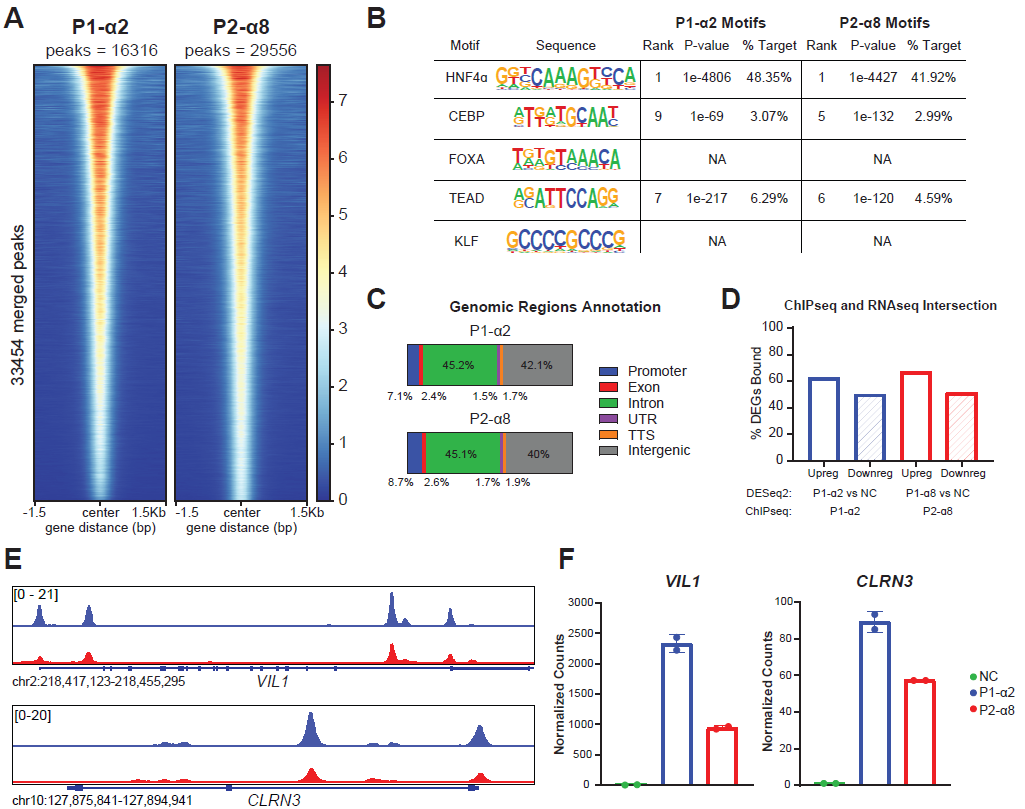

**Supplemental Figure S3: Exogenous P1-HNF4α have increased genomic binding at common target genes in Panc10.05.**

HNF4α ChIP-seq analysis following expression of P1-α2 (blue) or P2-α8 (red) in Panc10.05.

**A.** Heatmap showing occupancy of HNF4α at merged peak regions (significant peaks called by Macs2 in at least 1 condition). Replicates were combined for each isoform (n=2). Peaks ordered by descending mean signal across all datasets.

**B.** HOMER motif enrichment analysis for HNF4α, CEBPA, FOXA, TEAD, and KLF motifs.

**C.** Annotation of genomic regions bound by each isoform.

**D.** Percentage of significant DEGs (log2FC >.585 or <.585, padj <.05) bound by each isoform.

**E.** ChIP-seq tracks of HNF4α target genes *VIL1 and CLRN3* (combined replicates shown, n=2).

**F.** Normalized RNAseq counts of HNF4α target genes.

**SUPPLEMENTAL FIGURE S4**

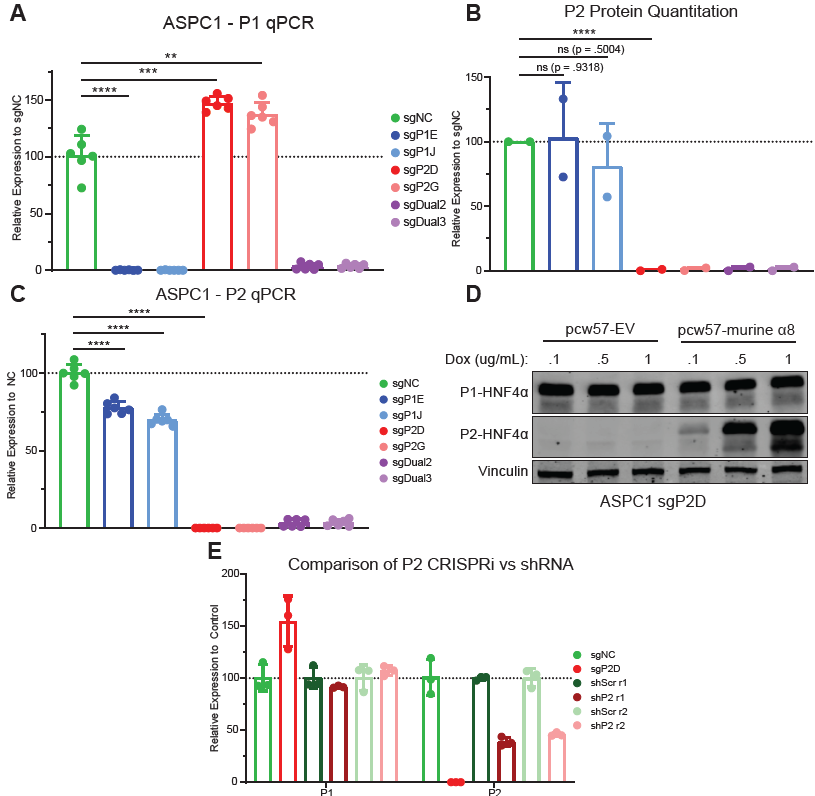

**Supplemental Figure S4. CRISPRi of P2 isoforms leads to an increase in P1 isoforms.**

**A.** qPCR of P1 RNA expression after CRISPRi in ASPC1, Student’s T-test, ** = p<.01, *** = p<.001, **** = p<.0001.

**B.** Quantitation of P2 protein expression in ASPC1 and HPAFII in **Figure 2A**. Signal intensity was normalized to β-Tubulin then to the sgNC of each cell line, Student’s T-test, ns = not significant, **** = p<.0001.

**C.** qPCR of P2 RNA expression after CRISPRi in ASPC1, Student’s T-test, **** = p<.0001.

**D.** Immunoblot of P1-HNF4α, P2-HNF4α, and Vinculin in P1-high cells (sgP2D) in ASPC1 after dox-induced re-expression of P2 (P2-α8).

**E.** qPCR of P1 and P2 RNA expression after shRNA knockdown of P2 in ASPC1-dCas9 cells, 2 replicates shown. sgNC and sgP2D were included as negative and positive controls, respectively.

**SUPPLEMENTAL FIGURE S5**

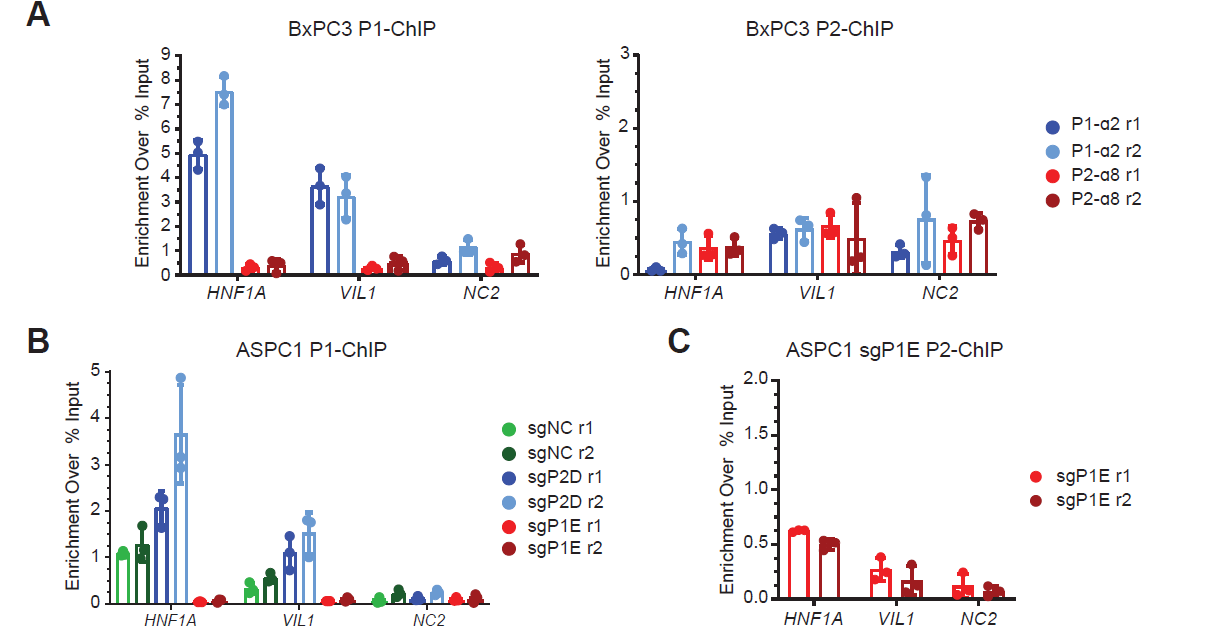

**Supplemental Figure S5. ChIP-seq using isoform-specific antibodies reveals increased P1 genomic binding after loss of P2 isoforms.**

Validation of isoform-specific antibodies for ChIP-seq in BxPC3 and ASPC1.

**A.** Left: P1 ChIP-qPCR of HNF4α targets after dox-induced expression of P1-α2 or P2-α8 in BxPC3. Right: P2 ChIP-qPCR of HNF4α targets after dox-induced expression of P1-α2 or P2-α8 in BxPC3. NC2 is a negative control region.

**B.** P1 ChIP-qPCR of HNF4α targets in the P1-high (sgP2D) or P2-high state (sgP1E) in ASPC1. NC2 is a negative control region.

**C.** P2 ChIP-qPCR of HNF4α targets in the P2-high state (sgP1E) in ASPC1. NC2 is a negative control region.

**SUPPLEMENTAL FIGURE S6**

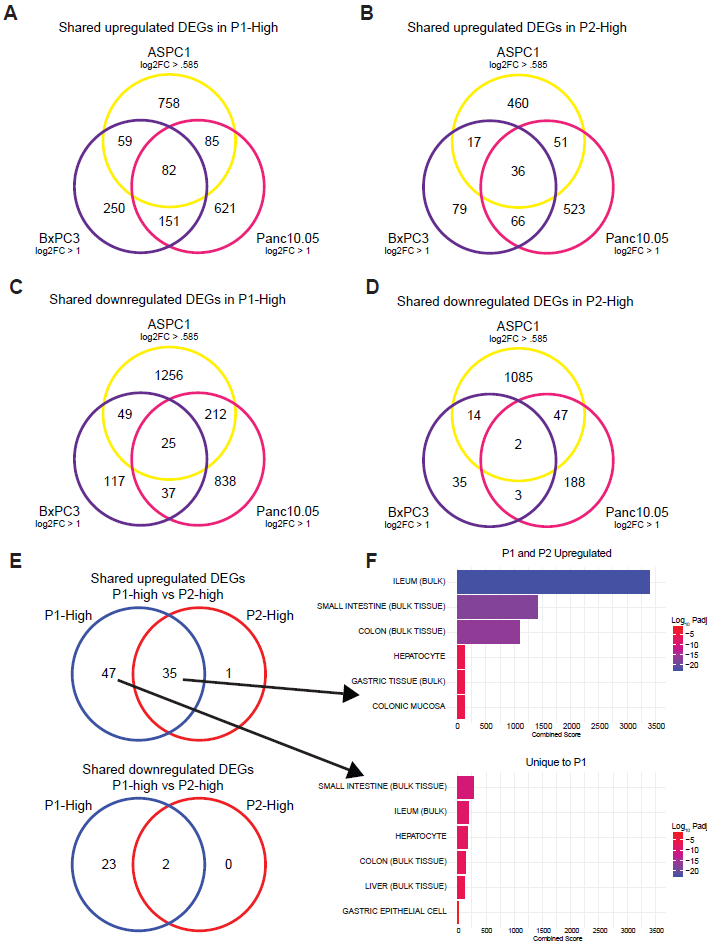

**Supplemental Figure S6. Integrated exogenous and endogenous differentially expressed genes.**

**A.** Venn diagram of DEGs significantly upregulated by P1 in BxPC3, Panc10.05 (log_2_FC > 1 or < -1 and padj < .05), and ASPC1 (log_2_FC > .585 or < -.585 and padj < .05).

**B.** Venn diagram of DEGs significantly upregulated by P2 in BxPC3, Panc10.05 (log_2_FC > 1 or < -1 and padj < .05), and ASPC1 (log_2_FC > .585 or < -.585 and padj < .05).

**C.** Venn diagram of DEGs significantly downregulated by P1 in BxPC3, Panc10.05 (log_2_FC > 1 or < -1 and padj < .05), and ASPC1 (log_2_FC > .585 or < -.585 and padj < .05).

**D.** Venn diagram of DEGs significantly downregulated by P2 in BxPC3, Panc10.05 (log_2_FC > 1 or < -1 and padj < .05), and ASPC1 (log_2_FC > .585 or < -.585 and padj < .05).

**E.** Venn diagram of consensus DEGs (significant in all three cell lines) regulated by P1 and/or P2. Above: upregulated; below: downregulated.

**F.** Enrichr analysis results for the indicated DEGs in the ARCHS4 Tissues gene set.

**SUPPLEMENTAL FIGURE S7**

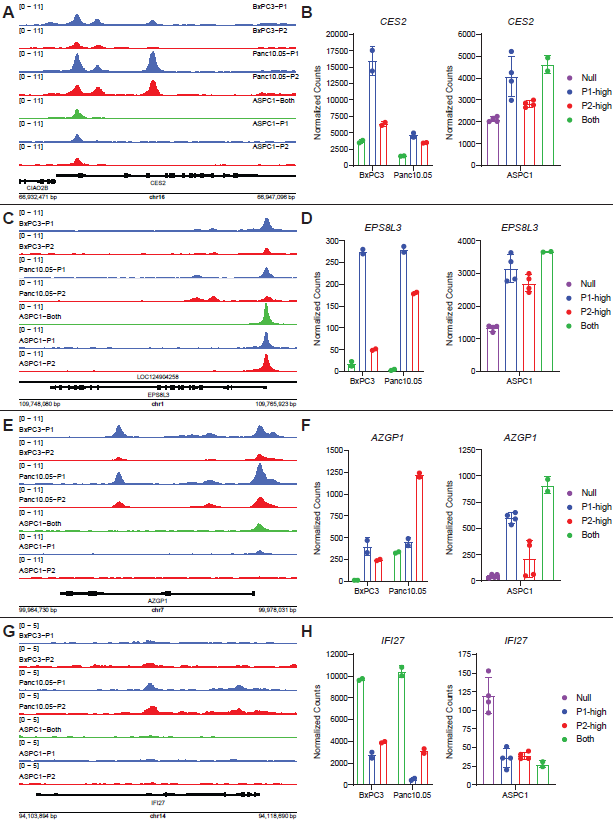

**Supplemental Figure S7. DNA binding and expression of shared target genes.**

Representative ChIP-seq tracks and normalized RNAseq counts, respectively, for DEGs upregulated by P1-only (**A and B**), upregulated by both isoforms **(C and D**), upregulated by P2-only (**E and F**), or downregulated by both isoforms (**G and H**).

**SUPPLEMENTAL FIGURE S8**

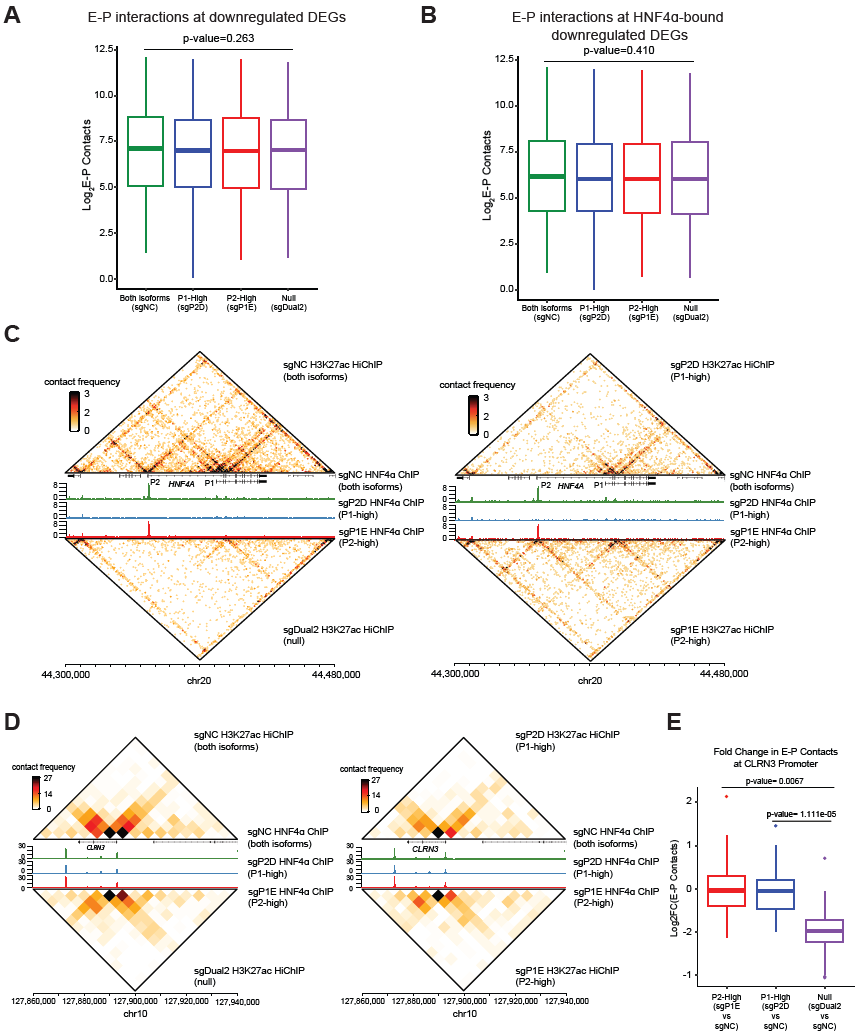

**Supplemental Figure S8. CRISPRi of HNF4α does not result in global changes in E-P contacts.**

**A.** Boxplot of enhancer-promoter contacts at genes downregulated in sgDual vs. sgNC RNAseq.

**B.** Boxplot of enhancer-promoter contacts at genes downregulated in sgDual vs. sgNC RNAseq and bound by HNF4α (sgNC HNF4α ChIPseq) either at promoters or the looped enhancers.

**C.** Heatmap of H3K27ac contact frequency with HNF4α ChIP peaks at the *HNF4A* locus. Left: sgNC (upper) and sgDual2 (lower). Right: sgP2D (Upper; P1-high) and sgP1E (lower; P2-high).

**D.** Heatmap of H3K27ac contact frequency with HNF4α ChIP peaks at the *CLRN3* locus. Left: sgNC (upper) and sgDual2 (lower). Right: sgP2D (Upper; P1-high) and sgP1E (lower; P2-high).

**E.** Boxplot of the fold change of E-P contacts at the *CLRN3* promoter in each contrast (HNF4α isoform knockdown vs. sgNC).

**Supplemental Tables**

**Supplemental Table S1.** Primary tumor and IPMN sample information and IHC quantitation, related to Figure 1.

**Supplemental Table S2.** Raw counts, normalized counts, and DEGs in BxPC3 and Panc10.05 cells expressing exogenous HNF4α isoforms, related to Figures 2 and S2.

**Supplemental Table S3.** Enriched gene sets by GSEA, DEGs in each cluster, and enriched signatures for each cluster by Enrichr in BxPC3 and Panc10.05 cells expressing exogenous HNF4α isoforms, related to Figures 2 and S2.

**Supplemental Table S4.** HNF4α ChIP-Seq peaks and gene annotations, and differentially bound HNF4α peaks and gene annotations in BxPC3 and Panc10.05 cells expressing exogenous HNF4α isoforms, related to Figures 3 and S3.

**Supplemental Table S5.** Raw counts, normalized counts, and DEGs in ASPC1 cells after CRISPRi of endogenous HNF4α isoforms, related to Figure 5.

**Supplemental Table S6.** Enriched gene sets by GSEA, DEGs in each cluster, and enriched signatures for each cluster by Enrichr in ASPC1 cells after CRISPRi of endogenous HNF4α isoforms, related to Figure 5.

**Supplemental Table S7.** HNF4α ChIP-Seq and P1 ChIP-Seq peaks and gene annotations, and differentially bound HNF4α peaks and gene annotations in ASPC1 cells after CRISPRi of endogenous HNF4α isoforms, related to Figures 6.

**Supplemental Table S8.** Genes significantly differentially expressed in all cell lines and Enrichr signatures, related to Figure S7.

**Supplemental Table S9.** Primers used in this study.
